# Threshold to brain quantum coherence

**DOI:** 10.64898/2026.10.14.759211

**Authors:** Philip P. Foster, Raj S. Chhikara, Aladin M. Boriek

**Affiliations:** Baylor College of Medicine, Department of Internal Medicine Mechanical Engineering, Houston, Texas, USA; Center for Space Medicine Mechanical Engineering, Houston, Texas, USA; Institute of Molecular Medicine, University of Texas Health McGovern Medical School Mechanical Engineering, Houston, Texas, USA; University of Houston-Clear Lake, Department of Mathematical, Applied, and Physical Sciences Mechanical Engineering, Houston, Texas, USA; Rice University, Mechanical Engineering, Houston, Texas, USA

## Abstract

Amongst brain structures, we identified a virtual void at the juxta– and paranode, within the intraperiod myelin layers, *V*_*IMI*_, as potential candidate for quantum near-ideal topological insulation. The insulation was extensively weighed here according to current findings. Values of biological and physical boundaries must be exact and a single missing/inexact element defining the anatomy, physiology, biochemistry, cytology, histology, and nano, leads to inaccurate boundaries/quantum equations of the myelin system. We determined the threshold of quantum coherence to decoherence. A propagating internodal electromagnetic interaction through the *V*_*IMI*_ void can be represented via Hamiltonian and entanglement. We tested three potential mechanisms for ion *K*^*+*^ data storage. 1. Quantum tunneling through a dielectric, *i.e*. single myelin layer. 2. Quantum tunneling through squeezing void, *V*_*IMI*_. The action potential is accompanied by mechanical axonal waves and myelin layers compression. Compression reduces the WKB action, and exponentially increases in transmission-tunneling probability. Resonance tunneling may add orders-of-magnitude boosts. 3. Classical squeezing passage. Quantum coherence, tunneling (1) and (2) are theoretically valid and testable, (1) at highly restricted parameter threshold. Unlike quantum tunneling, classical physic passage does not allow fine voltage-controlled metering, sorting, organizing the data storage, later rendering the retrieving impossible.

## A. Introduction

Understanding how information is encoded continues to elude current research efforts and the brain engram remains a knowledge gap. Intrigued by our hypothesis-driven statistical analyses of biological data, which suggested two distinct domains of investigation, a biological macroscopic measurements and quantum-scale mechanisms^1^. The equations of quantum mechanics are exact, and are exceptionally accurate models of reality that have passed every single experimental test for over a century. We had previously identified amongst brain structures a virtual void at the juxta− and paranode, within the intraperiod myelin layers, *V*_*IMI*_, as potential candidate for near-ideal topological insulation^1^; the proper insulation was extensively reviewed/weighed here according to current findings. Therefore, we split the brain and extensively inventoried its structures from anatomy down to quantum level as previously described^1^. Brain myelin compaction is a biological “vacuum-sealing” process which creates a tightly wrapped, highly efficient insulation around axons of white-matter tracts^2,3.^The myelin basic protein, MLP, “the molecular zipper”, neutralizes negative charges on membranes, pulling layers flatly together, in “compact myelin”, packing maximum insulation into a microscopic space^2,3.^If the brain cannot compact myelin properly, the action potential stops, as seen in neurodegenerative diseases (Alzheimer’s) or other demyelinating conditions.

Values of biological and physical boundaries must be exact, hereby, the equations of quantum mechanics are exact, and are exceptionally rigorous predictors of subatomic events in biology. A single missing/inexact element defining the anatomy, physiology, biochemistry, cytology, histology and nano, leads to inaccurate boundaries/quantum equations of the myelin system. We determined the threshold of quantum coherence to decoherence; as a theoretical model, it can be formulated as a quantum system under von Neumann equation. A propagating internodal electromagnetic interaction through the *V*_*IMI*_ void can be represented by an interaction Hamiltonian and entanglement is likely. We tested potential mechanisms for ion *K*^*+*^ data storage. 1. Quantum tunneling through a dielectric, *i*.*e*. single myelin layer. 2. Quantum tunneling through squeezing void, *V*_*IMI*_. The action potential is accompanied by mechanical axonal waves and myelin layers compression. Compression reduces the WKB action, producing an exponential increase in transmission-tunneling probability. 3. Classical squeezing passage. Quantum coherence, tunneling (1) and (2) are theoretically valid and testable, but at highly restricted parameter threshold. Classical physics would operate unrestrictedly; but unlike quantum tunneling, classical passage does not allow fine voltage-controlled metering, sorting, organizing the data storage, later rendering the retrieving impossible. The experimental verification of potential coherence may be forthcoming as teams are already reporting compressed ultrafast photography (CUP), of world’s fastest camera that achieves both an ultrafast imaging of up to 70 trillion frames per second (fps) and a large sequence depth of up to 1000 frames^4^. Today, data is efficiently stored in the cloud, remote servers of data centers, or in computers’ local hard drives. Hereby, we examined the available engineered models from AI computing, supporting data writing and storage which we compared with the brain myelin *V*_*IMI*_ system.

The organization of this paper follows a logical rationale. **A. Introduction. B. background. C. Results & Methods. Section C-I. Threshold of quantum coherence to decoherence**. i. Geometry of the intermyelinic intraperiod compartment volume V_IMI_… vi. The Hamiltonian for the two ensembles. xiv. Coherence versus entanglement. xv. Threshold… xvii. Separability. xvii. Conclusion. **Section C-II. Threshold to quantum tunneling. C-II-1. Two potential models of quantum tunneling. C-II-2. Quantum tunneling through a dielectric myelin layer**. i. Simulation of *K*^*+*^ crossing a dielectric. ii. Effective barrier height… v. *K*^*+*^ tunneling calculation. vi. Dielectric comparison myelin vs. phospholipid bilayer vs. 3D NAND flash. **C-II-3. Quantum tunneling through void, within the intraperiod compartment V**_**IMI**_**…** ii. *K*^*+*^ tunneling through the collapsing void. iii. Myelin layers’ compression. iv. WKB tunneling probability… vi. Tilted, time-dependent and collapsing barrier. vii. Sensitivity analysis, razor-edge threshold of parameters… ix. Compression inference… xviii. The moving barrier. xix. Resonant tunneling. **C-III. Classical passage. D. Discussion. D-I. Potential candidate for near-ideal topological insulation**… iii. Potential engram mechanism. **D-II. Coherence preservation**. i. Specific anatomy-histology preserving quantum coherence. ii. Intraperiod compartment. iii. Isolating from environment. iv. Unique physical properties. **D-III. Alzheimer’s disease provides further evidence. E. Conclusion**.

## B. Background

### B-I. *Transversal transfer* of charged ions across the axon, ion channels pumping *K*^***+***^ **and *Na***^***+***^

The architecture of the brain and AI-computers are both energy cost-driven and energy-savvy by design. The human brain achieves vastly superior energy efficiency, performing around 10^18^ mathematical operations (exascale equivalent, 1 quintillion) per second on just 15–20 watts of power, whereas computers consume about 20-40 megawatts to carry out the same 10^18^ operations per second. Transfer of information and energy are intertwined. In the brain, most energy is used to reverse ion influxes generating excitatory postsynaptic currents (EPSCs) and action potentials. EPSCs was restricted to minimize energy consumption, although kept at sufficient level not to jeopardize information transmission^5^. In the case of oligodendrocytes myelin sheath insulation, and Ranvier nodes, *qubit*_*Dep*_, may be virtual and initially created by voltage-gated ion channels and ion pumping. An electric current, *per se*, consists of electrons moving from one place to another in a conductor, usually in response to an electric field, which does not seem to be the case of the neuronal oligodendria-myelin-sheath insulation. Association of myelin sheath-Ranvier nodes provides remarkable insulation and rapid signal transmission in independent micro-circuitries packed in limited intra-cerebral space, hereby excluding shortcut commission errors. Energetic cell-induced node pumping generates an electric field from node to node. The original, elegant and concise mathematical heuristic model of transmembrane ionic transport at the node with the experiment backing the results was described by Hodgkin and Huxley^6^. Thereafter, much work was related to the *transverse*-ionic transport across membrane at the Ranvier Node^7^.

### BII. *Longitudinal jump* of electromagnetic potential along the axon, quantum-charged particles transferred from node to node

The *longitudinal*-*jump* from node to node remains elusive and conceals direct identification of the elemental vector of information in such micro-electromagnetic field. In quantum physics, this is also known as “*quantum teleportation*”. The physical particle does not move but rather transfers its exact quantum state (information) to another distant particle using entanglement. In physics, different supports exist, such as neural spin networks^8,9,^*J*-coupling^8,9,^electron spin^8,9,^spin-orbital coupling^8,9,^intra- and inter-molecular spin networks^8,9,^electron number (charge), photon, vibrational *qubit*, or others. Yet none of these theories has gained traction, although, in the universe of physics we are immersed in, there must be one elemental starter, yet to be discovered. Experience of protecting the brain in the physical space environment with absence of gravity and pressure led scientists to re-thinking the entire biology^10-12^.

## C. Results & Methods

### C-I. Threshold of quantum coherence to decoherence

#### i. Geometry of the intermyelinic intraperiod compartment volume *V*_***IMI***_

The inner cylinder has radius R, length L, and volume V_inner_,

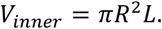

The outer cylinder has radius, R_outer_,

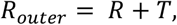

Therefore, the intraperiod space compartment volume, *V*_*IMI*_, is

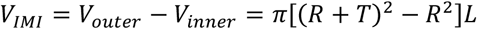

Simplifying,

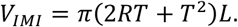

#### ii. Numerical parameterization with human brain biological values

The thickness of the intraperiod space compartment is

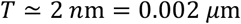

Hence,

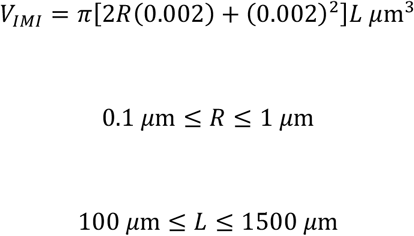

**Table 1.** Values of intraperiod space volumes, *V*_*IMI*_, for various axon lengths and radius.

| $R$ | $L$ | *intraperiod compartment*<br>*volume $V_{IMI}$* |
| --- | --- | --- |
| 0.1 $\mu\text{m}$ | 100 $\mu\text{m}$ | 0.1269 $\mu\text{m}^3$ |
| 0.1 $\mu\text{m}$ | 1500 $\mu\text{m}$ | 1.9038 $\mu\text{m}^3$ |
| 1 $\mu\text{m}$ | 100 $\mu\text{m}$ | 1.2579 $\mu\text{m}^3$ |
| 1 $\mu\text{m}$ | 1500 $\mu\text{m}$ | 18.8684 $\mu\text{m}^3$ |

*The boundaries are:*

Since

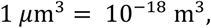

the volume of the intraperiod space compartment range is,

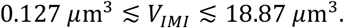

Or for further computations, in SI units,

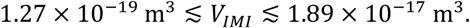

*Because* 1 *μm*^3^ = 1 femtoliter,

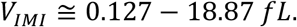

#### iii. *V*_*IMI*_ approximation

Because

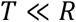

The term T^2^is very small. Hence,

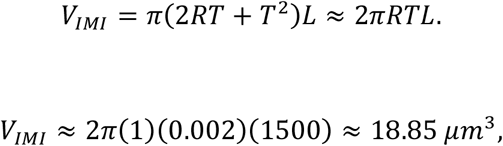

As illustrated in Fig. 2.

#### iv. Quantum description: density-matrix of the two ions ensembles

Now let’s formulate the quantum dynamics problem more precisely.

Let

- *S*_1_ = ensemble of approximately *N*_1_ = 10^7^ − 10^8^ *Na*^+^ ions,
- *S*_1_ = ensemble of approximately *N*_2_ = 10^3^ *Na*^+^ ions,
- *V*_*IMI*_ = the cylindrical volume within the void of the intraperiod compartment
- 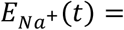 externally applied electric field driving S_1_,
- 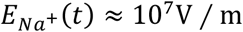 locally,
- *ρ*_12_ = density matrix of the combined two − ensemble system.

The complete quantum state is not commonly described by two independent density matrices. It is described by

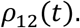

Acting here on the tensor product Hilbert space, for *Na*^*+*^ ions −note that in the next section, “Threshold to quantum tunneling”, the calculation applies to *K*^*+*^,

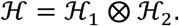

The time evolution of an isolated system is represented by the von Neumann equation

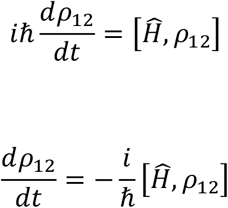

where

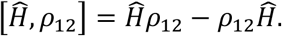

#### v. The Hamiltonian for the two ensembles

For the myelin system physiology, it is useful to separate the Hamiltonian (*Na*^*+*^ ions) into four contributions,

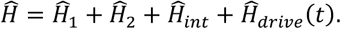

- Ĥ_1_ = describes ensemble *S*_1_,
- Ĥ_2_ = describes ensemble *S*_2_,
- Ĥ_*int*_ = describes the physical interaction between the two ensembles through the intervening region, intraperiod space compartment.
- Ĥ_*drive*_ = describes the physical interaction applied electromagnetic drive from *Node 1*.

### vi. The crucial distinction: interaction is not automatically entanglement

Initially, suppose the two ensembles are independent,

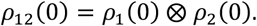

If the Hamiltonian (for *Na*^*+*^ ions) contains only local terms,

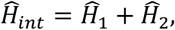

Then

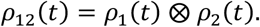

And the two systems remain separable. However, if there is a genuine coupling

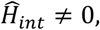

That simultaneously acts jointly on the two systems, the evolution can produce

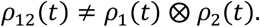

That is where entanglement can potentially arise.

But inequality alone is not the most rigorous definition of entanglement for mixed states. For a mixed density matrix, the correct question is whether *ρ*_12_ is separable,

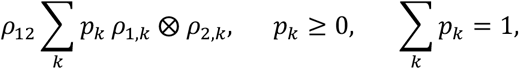

Hereby, the state is entangled.

#### vii. Significance of number of 10^7^ − 10^8^ *Na*^+^ ions in one node

This is an important concept. Ions *Na*^*+*^ in ensemble 1,

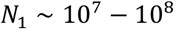

And ions *Na*^*+*^ (not yet activated) in ensemble 2,

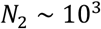

It would be too strong to say that all 10^7^ − 10^8^ *ions Na*^+^ are necessarily one giant entangled state. Instead, one should define collective degrees of freedom.

For example, if the relevant degree of freedom is spin, one can introduce a collective spin operator,

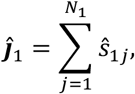

and,

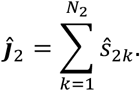

Then a possible effective interaction could have the form,

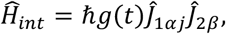

where *g*(*t*) represents the coupling strength and *α, β*, specify the relevant components.

This is an effective model, not a claim that your particular apparatus necessarily realizes this Hamiltonian.

#### viii. The electric field

We have the specific electric field, *E*_*Na*_,

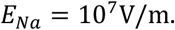

The electric interaction of a charged particle with an electric potential *ϕ* is represented by,

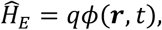

And for a *Na*^+^ion,

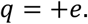

Therefore,

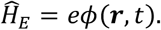

The important distinction is that an electric field propagating through the void can establish a physical electromagnetic coupling between the two regions. The coupling is not automatically a quantum-entangling interaction.

It can be:

1. classical electromagnetic communication,
2. quantum interaction,
3. or, a combination of both.

To establish entanglement, one needs to demonstrate that the resulting *ρ*_12_ *is nonseparable*.

#### ix The 100 mV amplitude change

The specific amplitude change at a node (e.g. Node 1) is approximately,

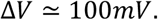

The corresponding electrostatic energy change of one singly charged *Na*^+^ is,

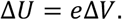

Since one electronvolt corresponds to one-volt times the elementary charge,

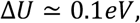

per *Na*^+^ ion for a 100 mV potential difference.

This is a useful quantity to distinguish from the electric field amplitude.

#### x. The “all-or-none” law

The specific triggering event follows the “all-or-none” law.

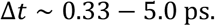

Distance between two nodes: 100 μm: Δ*t ∼* 0.33 ps for 1.5 mm Δ*t ∼* 5.0 ps. For n-impulse attempts (density), at Node 1, one reaches out the threshold to trigger the electromagnetic wave to reach Node 2. It approximately takes picoseconds for transmission between two nodes. Note that the AP velocity and this electromagnetic wave are totally different^1^.

#### xi. Zero-point motion does not destroy the quantum description

The statement about zero-point fluctuations is important.

A quantum particle does not generally possess an exactly fixed classical position. Its position is described by a quantum state or probability distribution.

For an idealized harmonic degree of freedom, the ground-state position uncertainty is

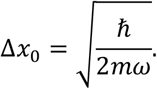

Thus zero-point motion is not evidence against quantum coherence. It is actually an intrinsic feature of quantum mechanics.

However, zero-point fluctuations are different concepts.

#### xii. The isolation assumptions

- No residual gas,
- No water or other atoms in the void,
- No phonons,
- No thermal fluctuations relevant to the isolated system,
- No fluctuating external electromagnetic field,
- No electrode/environmental degrees of freedom,
- Near ideal insulation surrounding the ensembles,
- No spontaneous electromagnetic emission relevant to the modeled system.

These assumptions will be discussed, one-by-one, and detailed later in the Discussion. This is the level where there is a threshold to quantum coherence and decoherence. Under those assumptions, the simplest theoretical description is indeed a closed-system unitary evolution,

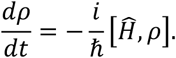

This is precisely the situation in which the von Neumann equation if appropriate.

But there is an important caveat: Isolation from the external environment does not by itself prove that the state is coherent. It merely removes or suppresses some important decoherence channels.

#### xiii. Coherence versus entanglement

These are related but different.

##### 1. Quantum coherence

A density matrix has coherence when it contains nonzero off-diagonal elements in the relevant basis,

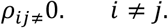

For example,

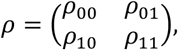

With

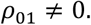

##### 2. Entanglement

Entanglement is stronger. It relates to the joint state,

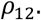

A state may have coherence without being entangled. Conversely, entanglement necessarily involves nonclassical correlations between subsystems, but its experimental identification requires an appropriate entanglement criterion or witness.

##### 3. Most useful equation for our system

For our proposed two-ensemble system (two-nodes), the formulation of the problem is,

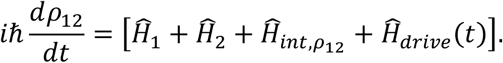

Initially,

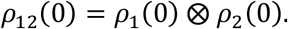

After the single activation,

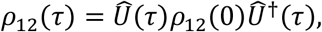

where

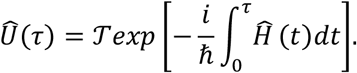

For a time-independent Hamiltonian, this simplifies to,

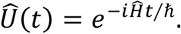

#### xiv. Threshold of quantum coherence to decoherence

*As a theoretical model, this can be formulated as a quantum system*. The appropriate mathematical description is the joint Hilbert space,

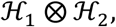

with joint density matrix

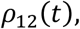

and von Neumann evolution

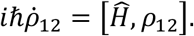

The assumptions of strong environmental isolation make this a reasonable close-system idealization. But there is a critical distinction,

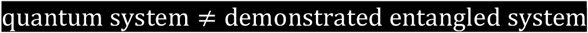

and also,

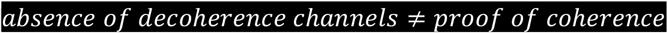

To claim experimentally that *S*_*1*_ and *S*_*2*_ become entangled, we would need to establish that the measured joint state satisfies an entanglement criterion.

#### xv. What would actually demonstrate entanglement?

Conceptually, we would need to establish something like

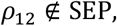

Where SEP is the set of separable states.

For a simple two-*qubi*t model, one could use concurrence, negativity, an entanglement witness, or a Bell-type measurement. For large ensembles such as *N*_1_ and *N*_2_, the appropriate observable and witness would depend strongly on what collective degree of freedom is actually controlled and measured.

A useful quantity is the partial transpose,

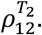

If the partial transpose has negative eigenvalue, the state is necessarily entangled.

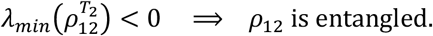

For sufficiently bipartite systems this gives the Peres-Hodorecki criterion; for large dimensional mixed systems, positivity of the partial transpose is not by itself sufficient to prove separability.

##### Note about this specific system

We can state rigorously,

Our proposed system can be seen as an isolated quantum system with two subsystems *S*_1_ and *S*_2_ evolving under von Neumann equation.

A propagating electromagnetic interaction through the void can be represented by an interaction Hamiltonian Ĥ_*int*_. If that interaction produces a nonseparable *ρ*_12_, then the two ensembles are quantum mechanically entangled..

But the values *N*_1_ *∼* 10^7^ − 10^8^, *N*_2_ *∼* 10^3^, *E*_*Na*_ *∼* 10^7^ V/m, Δ*V ∼* 100 mV, and a picosecond triggering time do not, by themselves, establish entanglement. They are parameters of the proposed physical model. The decisive quantity is the resulting joint density matrix and its experimentally demonstrated nonseparability. This question remains and we demonstrate the separability of *S*_*1*_ and *S*_*2*_ below.

#### xvi. The two ensembles S_1_ and S_2_ remain distinct and separable

*S*_*1*_ and *S*_*2*_ remain separable and can be treated as distinct systems rather than combined, *Node 1* and *Node 2* never merge. Their states and outputs are constantly identified separately^6^. They are separable and they are interacting, although this electromagnetic coupling during AP propagation is not strong enough to erase their distinction and separability. Ensembles *S*_*1*_ and *S*_*2*_ communicate within the compacted intraperiod compartment volume *V*_*IMI*_. However, there is no transfer of *Na*^*+*^ between the two systems, *S*_*1*_ and *S*_*2*_, across the void of *V*_*IMI*_. During the refractory period, Ensemble *S*_*1*_, of *Na*^*+*^, at *Node 1*, return to their basal resting state, after the passage of AP, while Ensemble *S*_*2*_ is now activated through the cycle we just described, and so on for the next steps and nodes. Mathematically, the overall system can still be described in terms of the two components

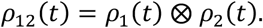

### C-II. Threshold to quantum tunneling

This is also critical on whether the system can work. The previous event, as described in Section C-I, involving *Na*^*+*^ is totally finished. Therefore, a totally different event involving *K*^*+*^ is now taking place. For *K*^*+*^ overshoot and 80 mV, potential difference^13^, the electric field (Fig. 1) across the cell membrane is

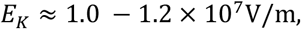

and the electric field *E*_*K*_ points inward toward the center of the axon. Note that there is only one role option, driven by the inward *Na*^*+*^ and outward *K*^*+*^ during impulse, according to the geo-localization of myelin layers, intraperiod space compartment, *V*_*IMI*_, axonal cell membrane at the paranode and juxtaparanode. In this entire paper, AP-inward-driven *Na*^*+*^ and AP-outward/driven *K*^*+*^ cannot have their respective roles interchanged. This postulate is valid for the AP and the potential quantum tunneling.

**Fig 1.**
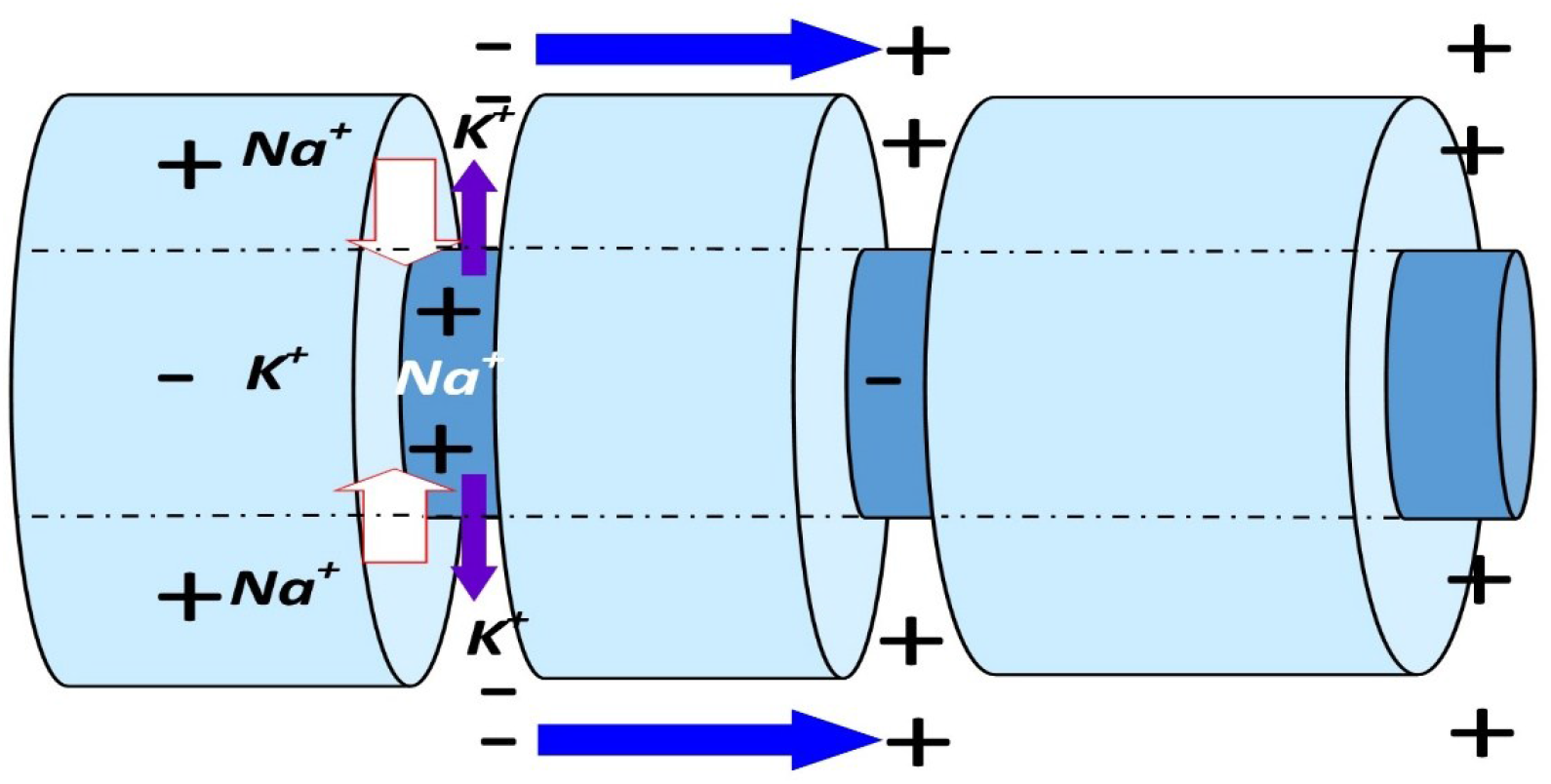
*Transversal transfer* across cellular membrane: ion channels *K*^*+*^/*Na*^*+*^ pumping (ATP-energy-system) driving ions against their electrochemical gradients. Excess of ions and net charges in a compartment are denoted by element-symbols (*K*^*+*^, *Na*^*+*^) and polarities (-, +). At rest: -70 mV relatively-negatively charged intraneuronal/axonal compartment. At depolarization, AP: +30-40 mV intraneuronal, whereas extraneuronal space becomes negatively polarized. *Longitudinal jump* of electromagnetic potential along the axon; negatively charged particles are transferred to the next node, aka *quantum teleportation* in quantum physics (marine-blue arrow). Myelin wraps: light blue cylinder; Axon: internal dark blue cylinder. Depolarization: *Na*^*+*^ is transferred intraneuronal (white arrow); *K*^*+*^ goes extracellular (purple arrow). The *NET* exchange results in negatively charging the extraneuronal space because more *Na+* are transported inside the axon. This phenomenon is important for understanding subsequent mechanisms underlying the engram formation.

**Fig 2.**
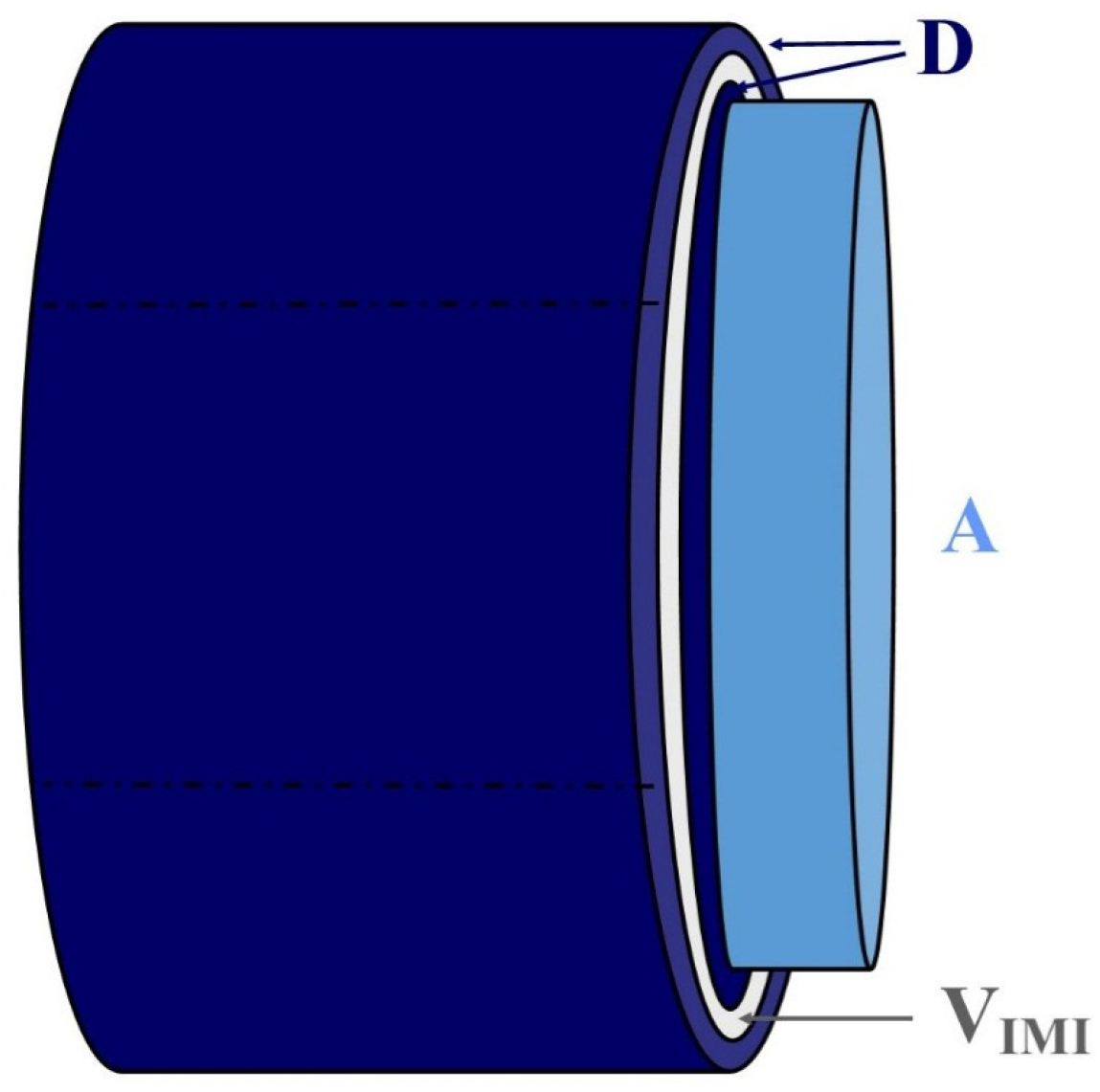
Geometry of one axon (A, pastel light-blue) and two myelin layers (D, dark blue). The intraperiod space compartment volume, *V*_*IMI*_, is in white (*V*_*IMI*_). We are calculating the real volume *V*_*IMI*_, the near-ideal topological quantum insulator (See one-by-one detail in Discussion). *V*_*IMI*_, is the compartment in which the quantum events are likely to occur during AP and *K*^*+*^ storage.

**Fig 3.**
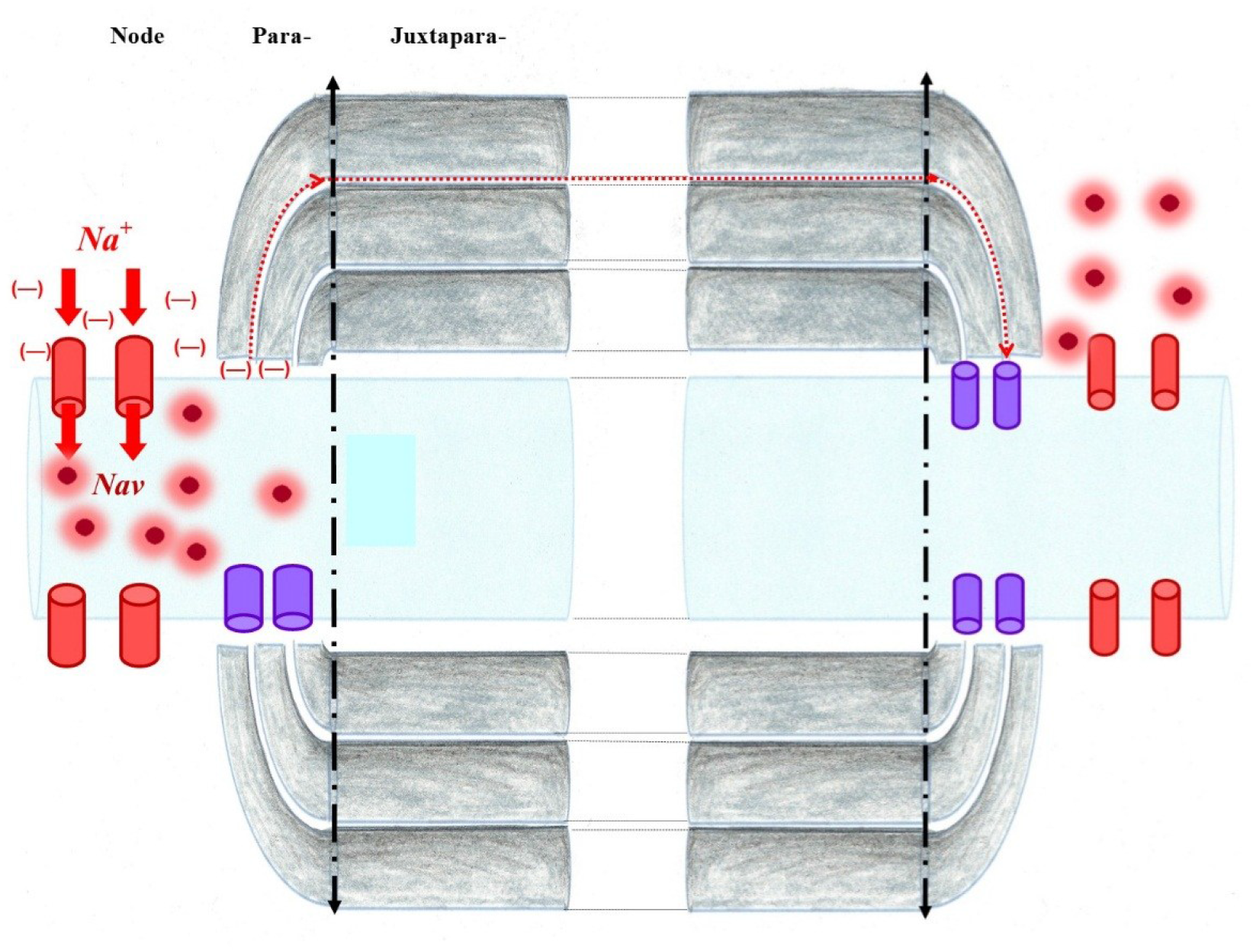
The ensemble *S*_1_ on the left, and *S*_2_. to the right. At its onset, the AP triggers the ion channels *Na*^*+*^ pumping driving *Na*^+^ ions inward within the axon (light blue), creating a potential difference [red (-)], Δ*V* ≃ 100*mV*, in the *S*_*1*_ region. Our postulate is that quantum interaction (dashed red arrow) between *S*_1_ & *S*_2_, takes place through the void of the intraperiod space, *V*_*IMI*_, offering near-ideal quantum topological insulation (See figures and details in Discussion). Ions Na+, are in red (quantum atomic model); extracellular depletion in the *S*_*1*_ region, *Node 1*.

**Fig 4.**
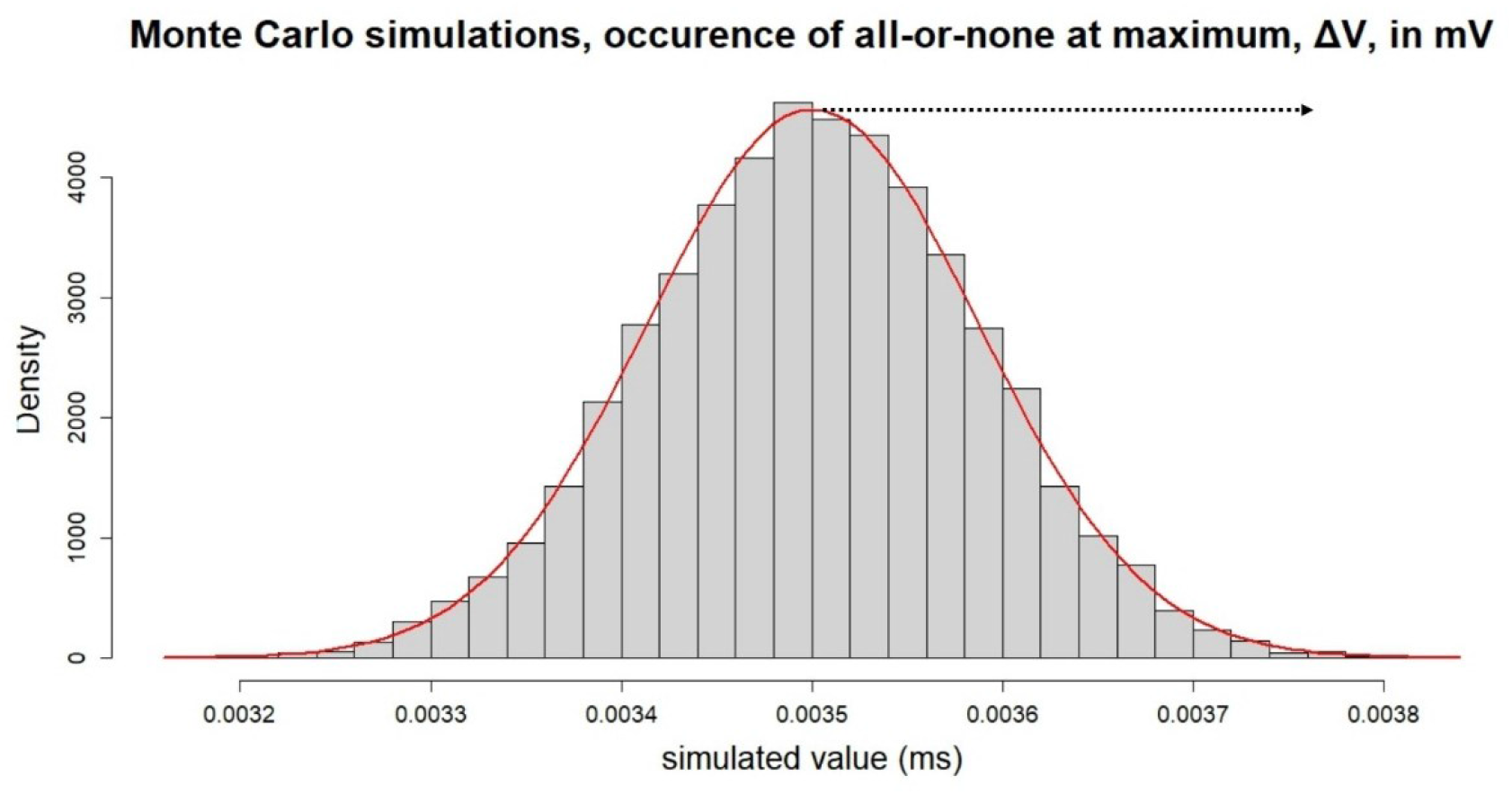
Monte Carlo simulation of the probability that one impulse out of a large number of impulses (density of impulses, Y-axis), may trigger a response, the “all-or-none” at a maximum, Δ*V*, likely Δ*V* ≃ 100 mV (dotted arrow). The single “all-or-none” response to a single sufficient impulse occurs and propagates the electromagnetic wave in the range of picoseconds, ps. The simulated values (X-axis) is the AP velocity ≈ 48 m/s. Note that this not a result. This is a demonstration to explain the composite nature of the electromagnetic signal in the brain: 1. Light speed component and 2. Slow component.

**Fig 5.**
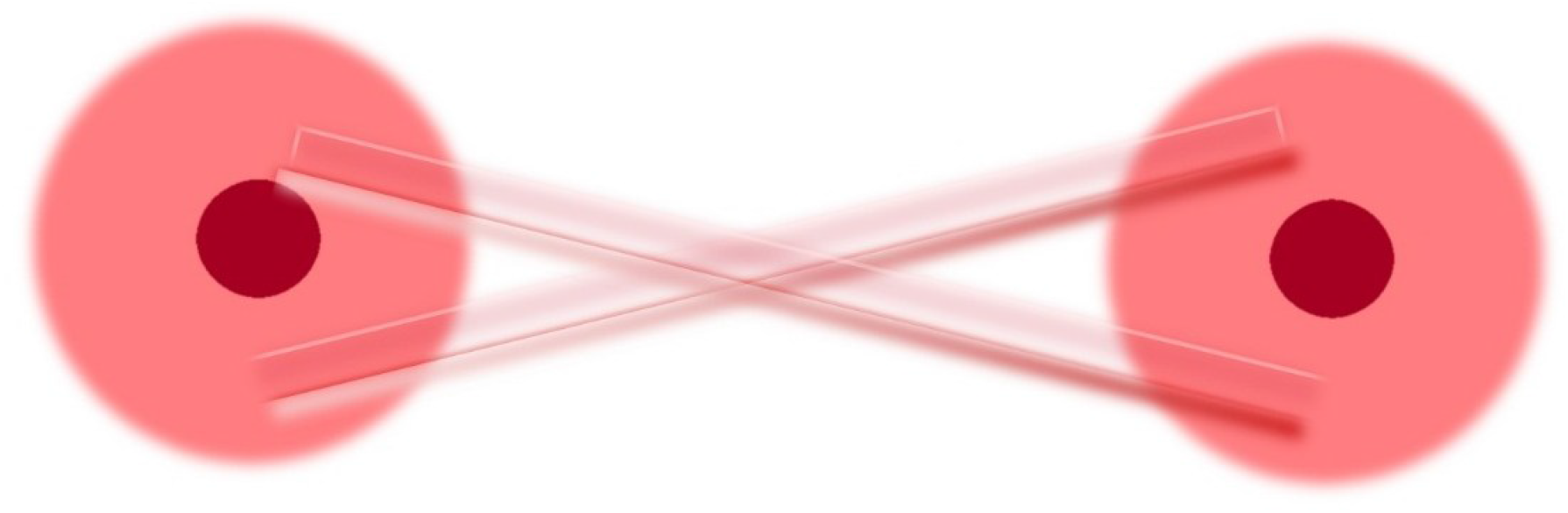
Two subatomic particles of ions *Na*^*+*^ become deeply interconnected, sharing a single unified quantum state regardless of the distance separating them.

**Fig 6.**
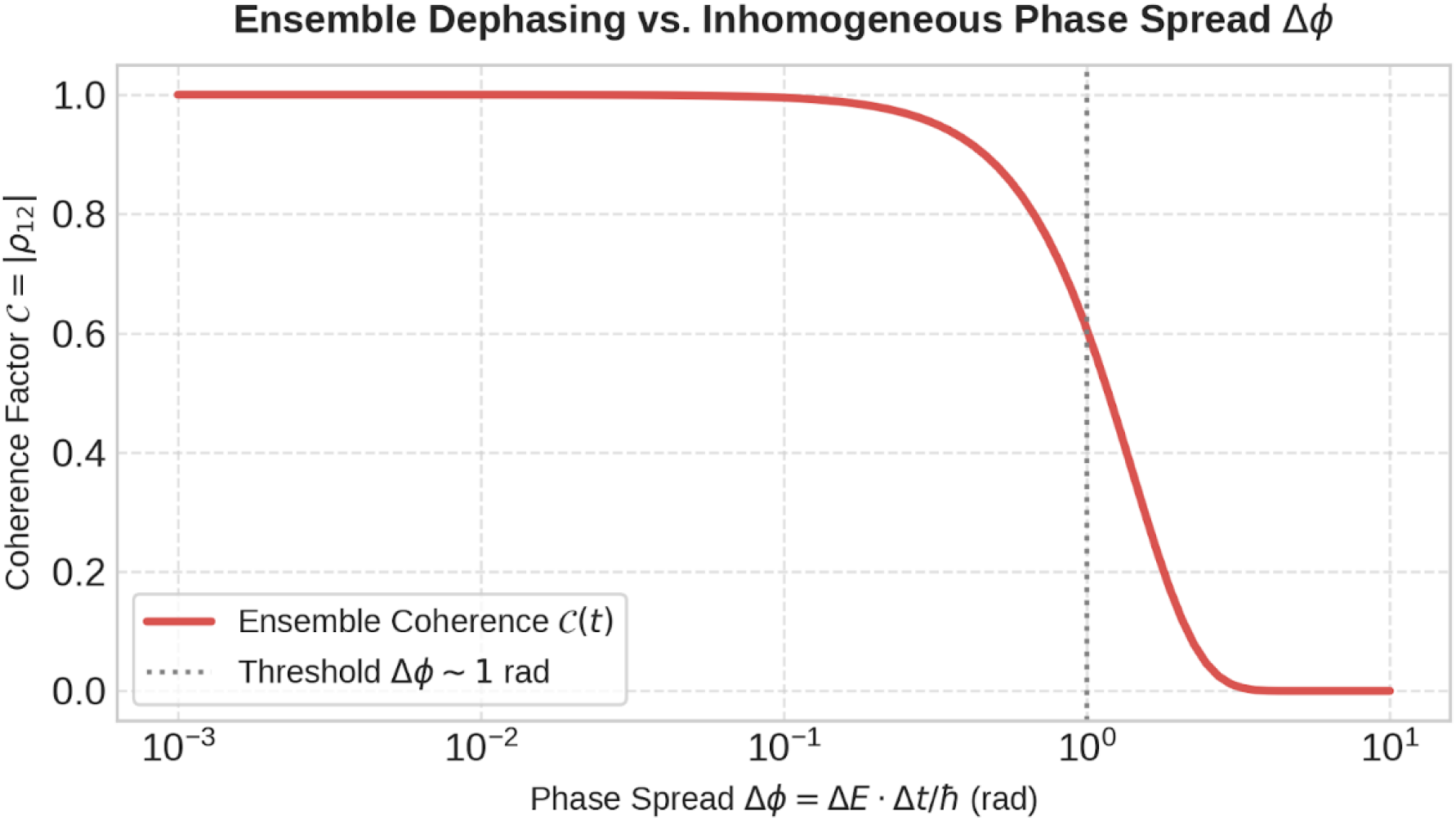
Threshold to decoherence. On the Y-Axis, the absolute value of the off-diagonal element of the two-level (or two-ensemble) density matrix, *C* = |*ρ*_12_|. It quantifies the remaining quantum coherence (phase relationship) between the relevant states after averaging over the ensembles. it is dimensionless and ranges 0 ≤ *C* ≤ 1. *C* = 1 is perfect coherence. *C* ⟶ 0, is complete loss of coherence. The curve shows how the coherence decays as the inhomogeneous phase spread 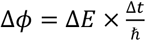 grows (dimensionless angle, expressed in radians). Where Δ*E* is the energy across the ions in the ensemble, Δ*t*, evolution time, and *ħ*, reduced Planck’s constant. Ions may accumulate different quantum phases even when no energy is exchanged with the external environment. This phenomenon is called inhomogeneous dephasing when the phase differences arise from a distribution of static/quasi-static frequencies or energies. Δ*ϕ* = 10^0^ = 1 rad, (dotted line) is the coherence threshold to decoherence. When Δ*ϕ* ≥ 1 rad, the different members of the ensemble acquire sufficiently different phases that their average coherence collapses meeting a sharp drop of the red curve.

**Fig 7.**
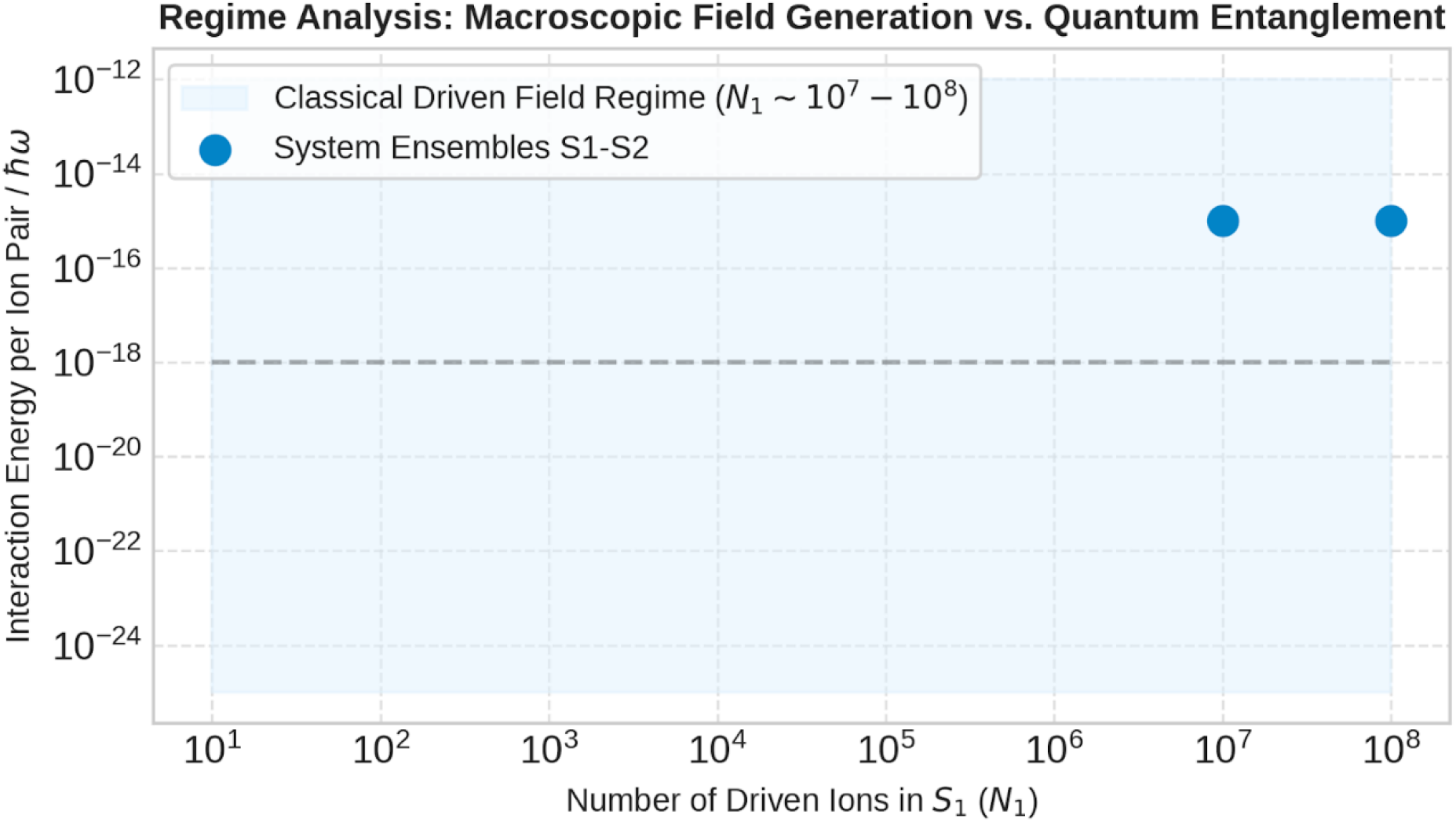
Quantum entanglement. Interaction energy per ion pair (Y-axis), vs. number of ions driven in Ensemble *S*_*1*_, at *Node 1*. In *S*_*1*_, one ion *Na*^*+*^ is pumped inward through the ion channels *Na*^*+*^, then swallowed and vanishing inward, leaving a negative difference of potential (**-**), at instant (t), creates an interaction energy (within picosecond if electromagnetic transmission alone, or instantaneously, if entangled, see below) with one ion *Na*^*+*^ freely floating in *S*_*2*_ (see Fig. 2). This interaction energy (dimensionless) is represented on the Y-axis, where *ω* is a characteristic frequency of the driven system (spin precession, Rabi frequency, etc.). The horizontal dashed line at 10^−18^ is a conventional scale separating “macroscopically classical field” behaviour from regimes where few particle quantum correlations might still be visible. The two blue points sit at *N*_1_ *∼* 10^7^ − 10^8^ (range of ions *Na*^*+*^), and lie well above the line, indicating the collective field generated by the large driven ensemble *S*_*1*_ is expected to act classically on the much smaller ensemble *S*_*2*_.

**Fig 8.**
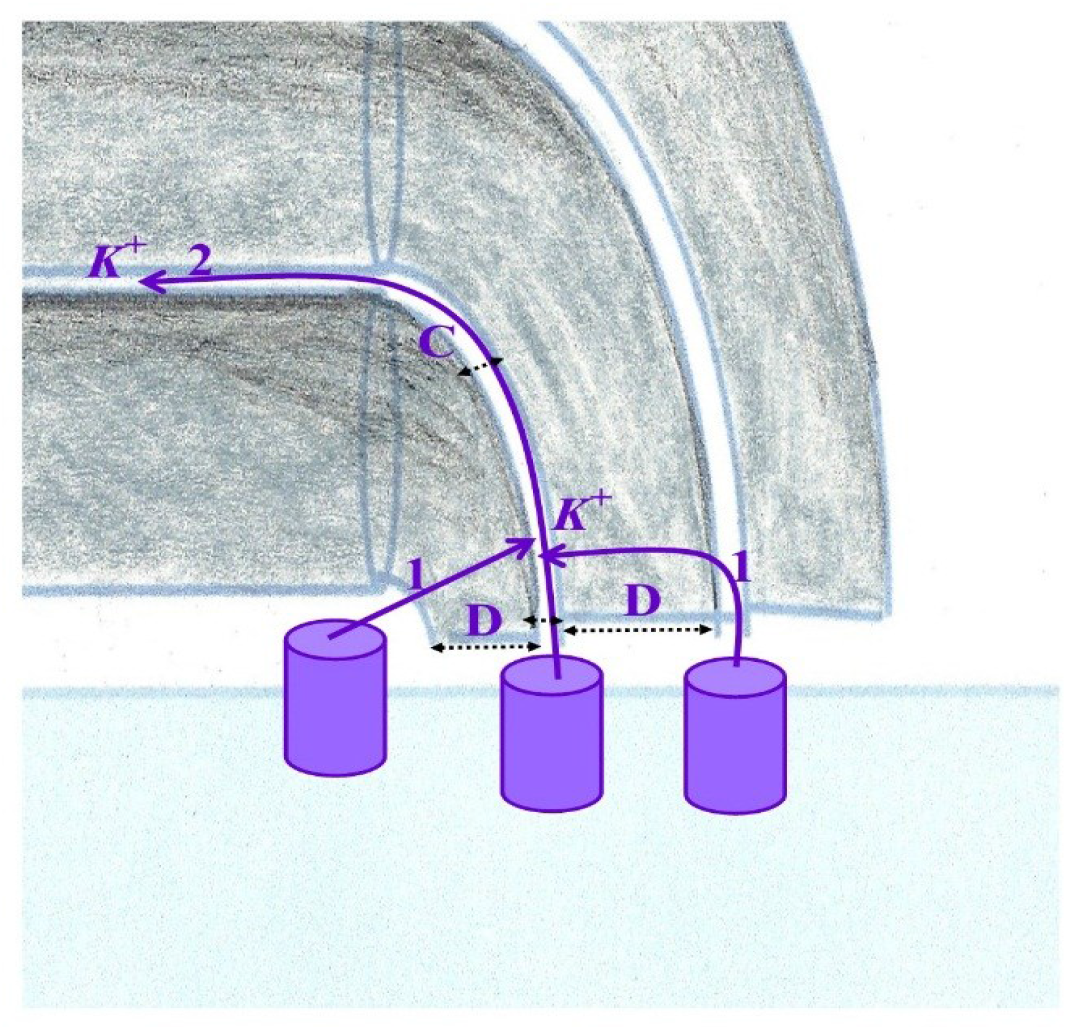
Two quantum tunneling option in the near-topological quantum insulation system, myelin intraperiod compartment, *V*_*IMI*_. 1). Across a single dielectric myelin layer (D), Model 1; and 2). Across the compressed void (D, initially dielectric, later charge-trap, C), of the myelin intraperiod compartment, *V*_*IMI*_.

**Fig 9.**
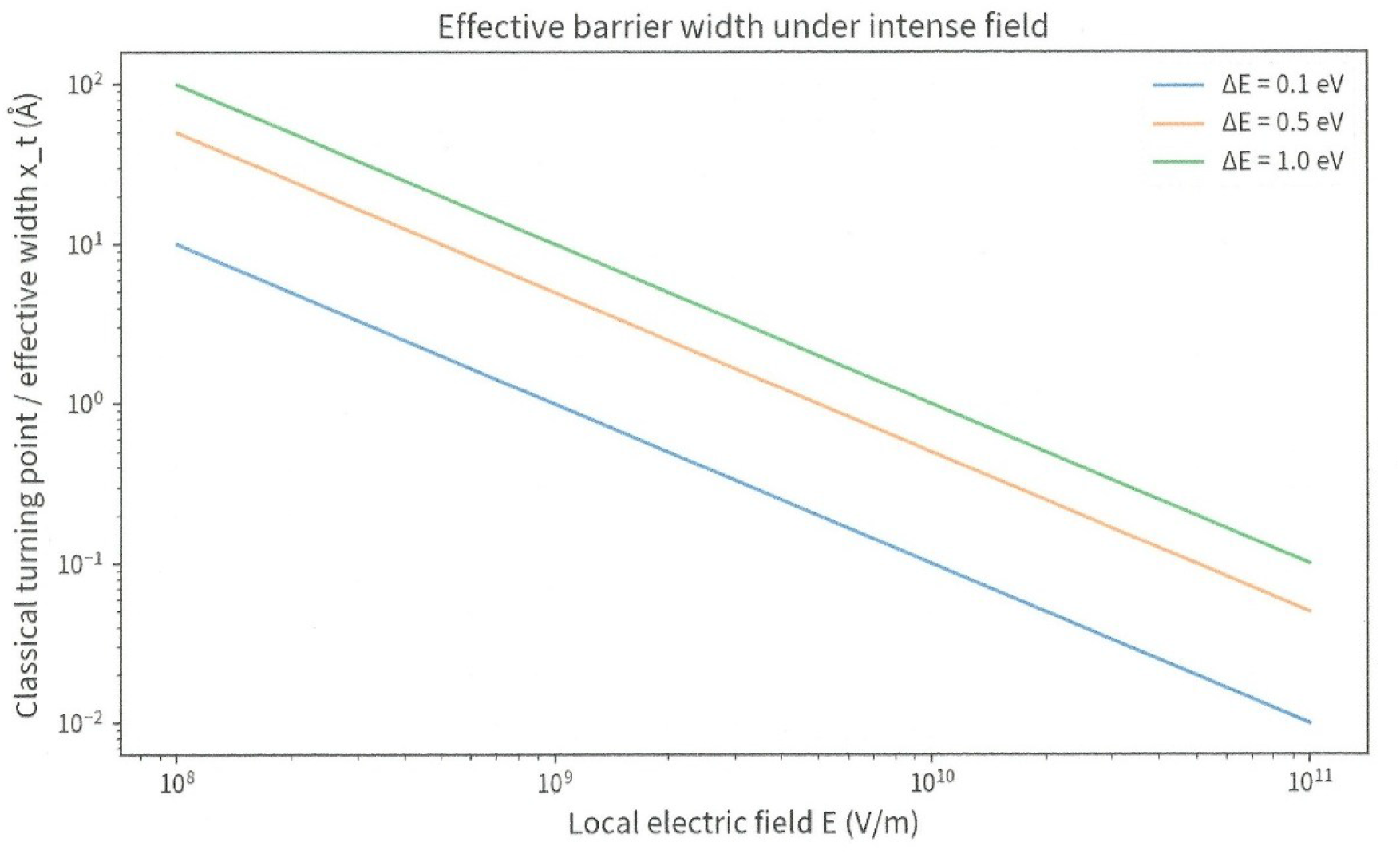
Effective barrier height as a function of the local electric field.

The geometry of this near-ideal topological quantum insulator, only allows a few available options for *K*^*+*^ storage exist within the intraperiod compartment. The questions are,

1. Would *K*^*+*^ quantum tunneling occur across one dielectric layer? The mass of the ion *K*^+^ is ∼7.1 × 10^4^ times that of the electron and its radius *r*_*K*_ ≈ 0.138 nm, are extremely limiting through the effective barrier height such as a single myelin layer.
2. Would *K*^*+*^ quantum tunneling occur through void, within the intermyelinic intraperiod compartment volume *V*_*IMI*_ ?
3. Would there be a classical passage of *K*^*+*^ through void, within the intermyelinic intraperiod compartment volume *V*_*IMI*_ ? (See next Section).

This is an important distinction, a *K*^*+*^ crossing an actual open void is not automatically a tunneling event. Whether it is tunneling depends on the time-dependent potential *V*(*x, t*) experienced by the ion while the gap closes.

### C-II-1. The two quantum tunneling models are genuinely different

#### 1. Dielectric barrier. Model 1

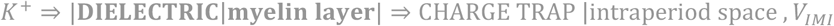

The inter myelin layer, *aka* intraperiod space compartment volume, *V*_*IMI*_, is the charge trap

#### 2. Compressing void. Model 2

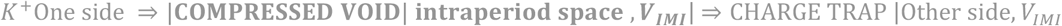

while *d*(*t*) ↓. Here, the important physics is the time-dependent confinement and interface potential.

### C-II-2. Quantum tunneling through the dielectric single myelin layer

#### i. Simulation of *K*^*+*^ crossing a dielectric, 3. 9 − 6. 8 nm, proximal myelin layer

This is where the new calculation become becomes particularly interesting. The dielectric/barrier thickness, proximal to the axon is

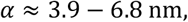

and the thickness, *T*, of the intraperiod space compartment, *V*_*IMI*_, *aka* the charge-trap region, is

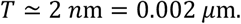

For a singly charged ion potassium,

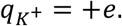

The electric field lowers/tilts the potential energy barrier, Δ*U*, by

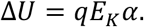

Because *q* = *e*, it is especially convenient to express this in electronvolts, unit of energy,

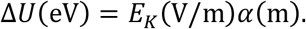

For *E*_*K*_ = 1.1 × 10^7^V/m, and *α* ≈ 3.9 nm,

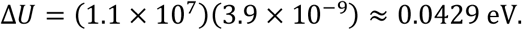

For *E*_*K*_ = 1.1 × 10^7^V/m, and *α* ≈ 6.8 nm,

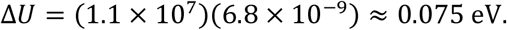

Thus,

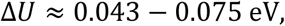

for the 3.9 − 6.8 nm dielectric barrier at 80 mV. This result is already useful as the electric field *E*, itself, contributes a potential-energy drop of approximately 43 − 75 meV, across the dielectric.

#### ii. Effective barrier height

For the quantum calculation, with a dielectric single myelin layer, the missing parameter is essentially

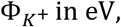

Or equivalently, for transfer into the hydrophobic dielectric from the channel environment of the dielectric system (totally unrelated to the *K*^*+*^ axonal cell membrane *Kv1* ion-channels)

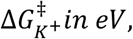

where 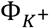 is the effective potential-energy barrier seen by *K*^*+*^ relative to its incidence/transmitting state. But there is an additional complication for *K*^*+*^ that did not arise in the same way in a simple electron-tunneling calculation. *K*^*+*^ is an ion moving in a biological, dielectric environment. Therefore, the relevant barrier is not simply the electron work function or a generic semiconductor band gap. We need to know the effective free-energy/potential landscape for *K*^*+*^.

#### iii. Effective ion *K*^*+*^ mass

The WKB tunneling exponent contains the effective ion *K*^*+*^ mass *m*_*k*_ ≈ 39.10 u. We should explicitly label that as an approximation, because the effective quantum dynamics of an ion in a condensed biological medium are not necessarily equivalent to a free particle with its vacuum mass.

#### iv. Barrier profile

Most importantly, we need to know whether the barrier is approximately rectangular or tilted. Because the electric field is intense, the most appropriate model is a trapezoidal/tilted barrier,

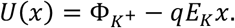

For *K*^*+*^

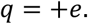

This is more physically appropriate than simply assuming a constant 0.10 − eV barrier.

#### v. *K*^*+*^ tunneling calculation

The electric field and 100-mV potential drop do not by themselves determine the tunneling probability. For quantum tunneling, we need the effective potential barrier *U*(*x*), seen by *K*^*+*^ relative to *K*^*+*^ energy. The rigorous calculation is consequently a parameterized approximate WKB (Wentzel-Kramers-Brillouin), transmission/tunneling probability

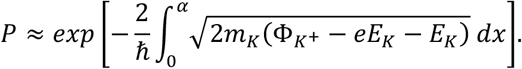

For a rectangular barrier, the WKB transmission probability is approximately

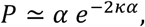

where

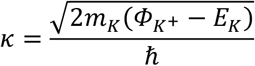

*α* ≈ 3.9 − 6.8 nm: is the dielectric thickness *m*_*K*_: is the *K*^*+*^ mass, *m*_*k*_ ≃ 39.10 u ≃ 6.49 × 10^−26^kg.

*U*(*x*) − *E*_*K*_: is the effective barrier energy, *aka* microscopic barrier height

*ħ*: is the reduced Planck constant.

#### vi. Dielectric comparison myelin *vs*. phospholipid bilayer *vs*. 3D NAND flash

The myelin composition and biophysical properties are also critical. This is the most important correction to the previous calculation. The eV barrier is insufficient to demonstrate the testability. A single myelin layer, proximal to the axon, is much closer in low-*κ* dielectric environment of a semiconductor than the high-*κ* charge-trap nitride layer of a 3D NAND cell. Figure 10 was derived from recent studies^14-17^. However, it is not equivalent to a specific NAND material. For quantum tunneling calculations, the closest useful analogue is generally the low-*κ* SiO_2_ tunnel dielectric, provided the comparison is made to the lipid rich hydrophobic myelin. Quantum tunneling may only occur within the first or second myelin layers proximal to the axon. Addition of myelin wraps outward of the axon provides further beneficial near-ideal topological insulation preserving quantum coherence of the system. However, piling myelin layers alters the quantum tunneling ability at a distance away from the axon which does not matter to the system since it is not a useful effect. It may only limit the storage capacity to the proximal myelin layers.

1. The 2026-2027 3D NAND charge-trap cell is not one dielectric. A representative 3D NAND stack contains a blocking dielectric, a charge-trap, and a tunnel. One published 3D NAND model uses AI_2_O_3_(*ε*_*r*_ ≈ 5.7), Si_3_N_4_(*ε*_*r*_ ≈ 9.7), and SiO_2_(*ε*_*r*_ ≈ 4.2), respectively, with a 5-nm SiO_2_ tunnel layer. The exact stack varies by manufacturer and generation.
2. Myelin composition. Lipid/protein ratio: 80/20% (standard cell membrane ratio: 40/50%). Myelin is uniquely enriched in galactocerebrosides (GalC), sulfatides, plasmalogens, cholesterol (∼40% total lipid mass), which packs tightly to prevent water penetration. Myelin proteins provide structural adhesion, myelin basic protein (MBP), positively charged peripheral protein that electrostatically seals the negatively charged inner cytoplasmic faces together into the major dense line (MDL). The proteolipid protein (PLP), is a transmembrane protein which zips opposing extra-axonal myelin layers together (compacting) into the intraperiod lin (IPL), aka intraperiod compartment volume, which we designed as ***V***_***IMI***_, in our computations. ***V***_***IMI***_, is a virtual hydrophobic void, in which water was squeezed out. The outward further centripetal layers potentially provide near-ideal topological insulation; this will be discussed in the Discussion section.
3. The dielectric constant is not dielectric potential; this is an important correction. The dielectric constant *ε*_*r*_ is a material property. The electric potential V, electric field E, and quantum tunneling depend on geometry, barrier thickness, barrier height, charge, applied voltage, boundary conditions, and dielectric distribution, as described above. The one-myelin proximal layer (*ε*_*r*_ ≈ 3.02) is close to SiO_2_(*ε*_*r*_ ≈ 4.2), 3D NAND dielectric, and closer than a phospholipid bilayer (*ε*_*r*_ ≈ 2.5), such as the axonal cell membrane. Likely making the myelin layer a better candidate than the cell membrane. In addition, it is more rigorous to say that myelin (*ε*_*r*_ ≈ 3.02) is closer to the low-*κ* dielectric materials, SiO_2_(*ε*_*r*_ ≈ 4.2), of the 3D NAND dielectric, than to the high-*κ* charge-trap Si_3_N_4_(*ε*_*r*_ ≈ 9.7) layer, used as charge-trap. This does not affect our system since intraperiod space compartment, *V*_*IMI*_, *aka* the charge-trap region, is virtual void.

**Fig 10.**
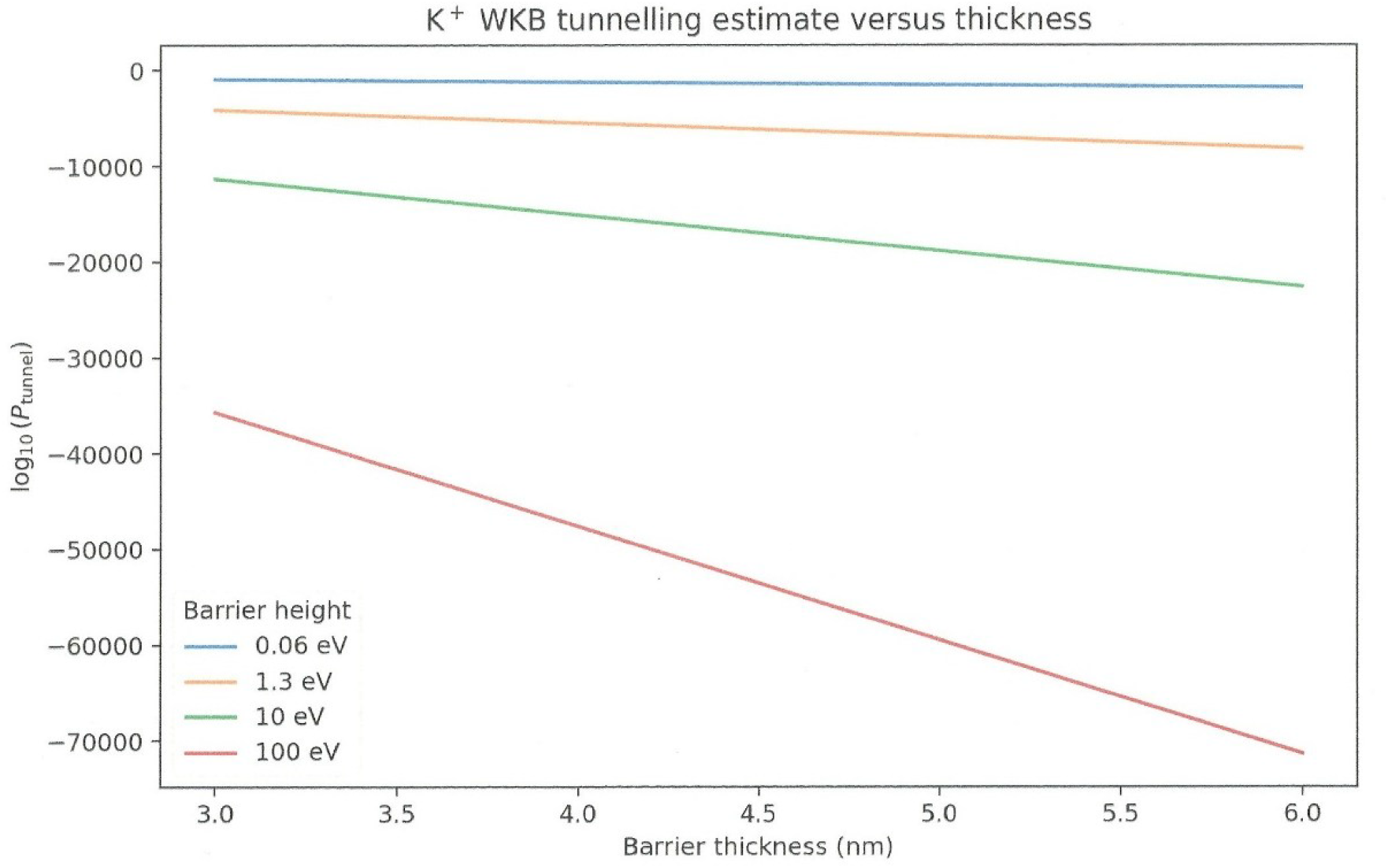
Transmission probability P of quantum tunneling. On the Y-axis, log_10_(*P*). When log_10_(*P*) ⟶ 0, top-left of the graph, the probability of quantum tunneling increases, when log_10_(*P*) = −0.1, then *P* ≈ 0.79.

**Fig 11.**
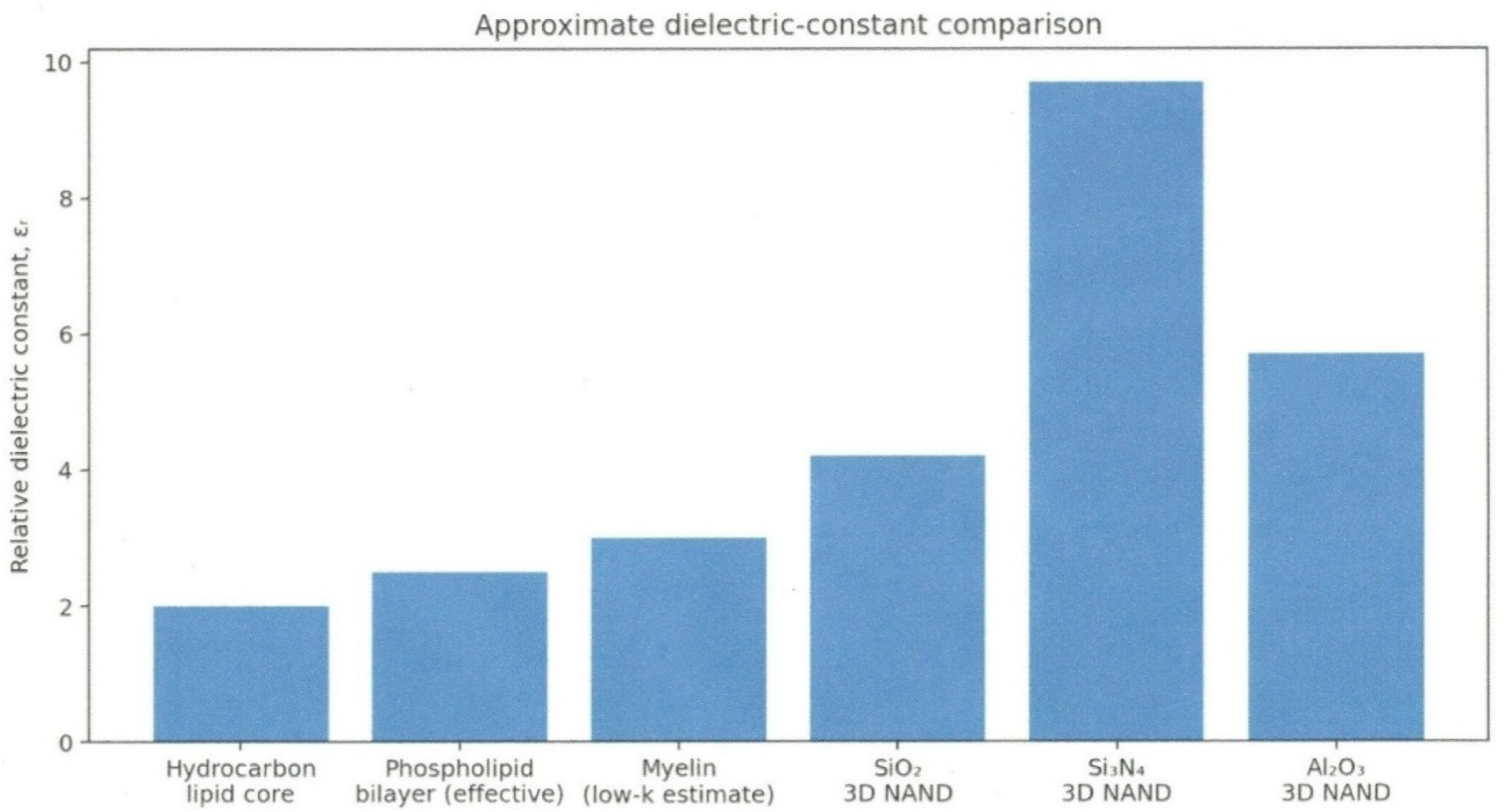
Dielectric constants of various material. The single myelin layer constant (*ε*_*r*_ ≈ 3.02), is the closest to the dielectric SiO_2_(*ε*_*r*_ ≈ 4.2) of the 3D NAND. Note that Si_3_N_4_(*ε*_*r*_ ≈ 9.7), is used as charge-trap layer.

Quantum tunneling is the fundamental physical mechanism as the gateway that enables the movement of charges (electrons or positive charges), in 3D NAND flash memory^18-20^. Quantum tunneling is organizing individual charges into separate bins like sorting mail, 3D NAND quantum tunneling “sorts” data by metering the exact quantity of charges and electrons allowed to pass through a quantum barrier^18-20^. It also depends on the physical properties of the charge-trap layer and precise voltage controls. If no quantum tunneling exists in the system; there is absence of memory, and further structured and sorting becomes impossible.

In the 3D NAND, silicon-nitride, Si_3_N_4_, the charge-trap, is separated from the charges (electron) source by an incredibly thin insulating dielectric barrier SiO_2_. When the precise voltage is applied at the control gate, it creates an intense electric field, E. This field tilts the energy bands of the insulator (dielectric), allowing charges (electrons) to undergo Fowler-Nordheim (FN) quantum tunneling−”effectively teleporting” straight through the energy barrier into the charge trap space. Thereafter, the silicon-nitride, Si_3_N_4_, the charge-trap, acts as a microscopic sponge filled with thousands of localized atomic alveoli (“defects” or “traps”), which completes a sorting by dispersed and complete immobilization. Another parameter, the threshold voltage, precisely controls the volume of charges (electrons) pushed through the quantum barrier. This process occurs by the memory controller applying incremental pulse programming^21^.

#### vii. Magnetic field

A magnetic field can be measured outside the skull *via* magneto-encephalography (MEG), but it does not assess deep in the brain directly within the myelin system coated by its near-ideal topological insulation with additional several white matter tracts piling up (Fig. 6). In a near future, techniques such as neuropixels^22,23^may enable to directly measure the magnetic field within the myelin system. Hereby, no accurate magnetic field measurement of the intraperiod compartment is available; therefore, the present model considers the measured electrostatic potential and does not introduce an experimentally unsupported magnetic field term. Note that a magnetic field term may help advocating and defend the case of this hypothesis.

#### viii. Partial conclusion of this option

The proposed *K*^*+*^ quantum tunneling is experimentally testable. Based on biological data, the present calculations define quantitative predictions that can be compared with measurements of *K*^*+*^ transfer as a function of dielectric thickness and electric field. Although the equations of quantum mechanics are exact, they are exceptionally accurate models of reality, our assessment must also pass the experimental test for full validation.

### C-II-3. Quantum tunneling through the intraperiod compartment *V*_***IMI***_

#### i. Modeling through void, within the intermyelinic intraperiod compartment volume *V*_***IMI***_

We must reformulate a new geometry.

**insulator 1**, myelin layer **1** | **time** − **department void *d***(***t***) | **insulator 2**, myelin layer **2**
with

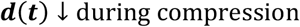

and a *K*^*+*^ ion approaching the gap (Fig. 12). The important variable is not simply a fixed barrier width d, but

**Fig 12.**
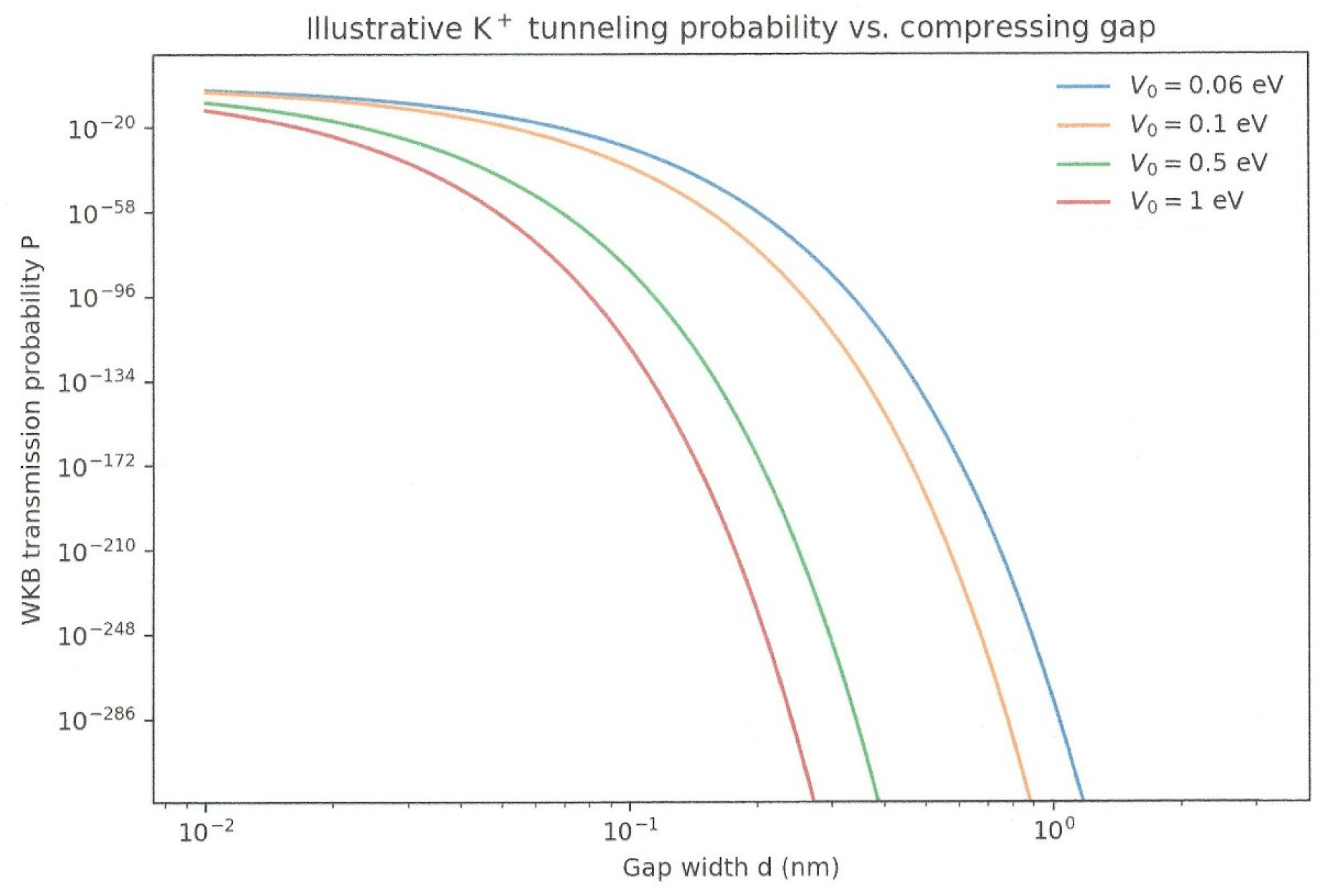
Tunneling probability of *K*^*+*^ *vs*. initial compressive gap. When the adjacent layers are flatly pulled together into “compact myelin”, the quantum tunneling probability increases for an initial gap in the range of ∼0.1 nm, under WKB alone.

**Fig 13.**
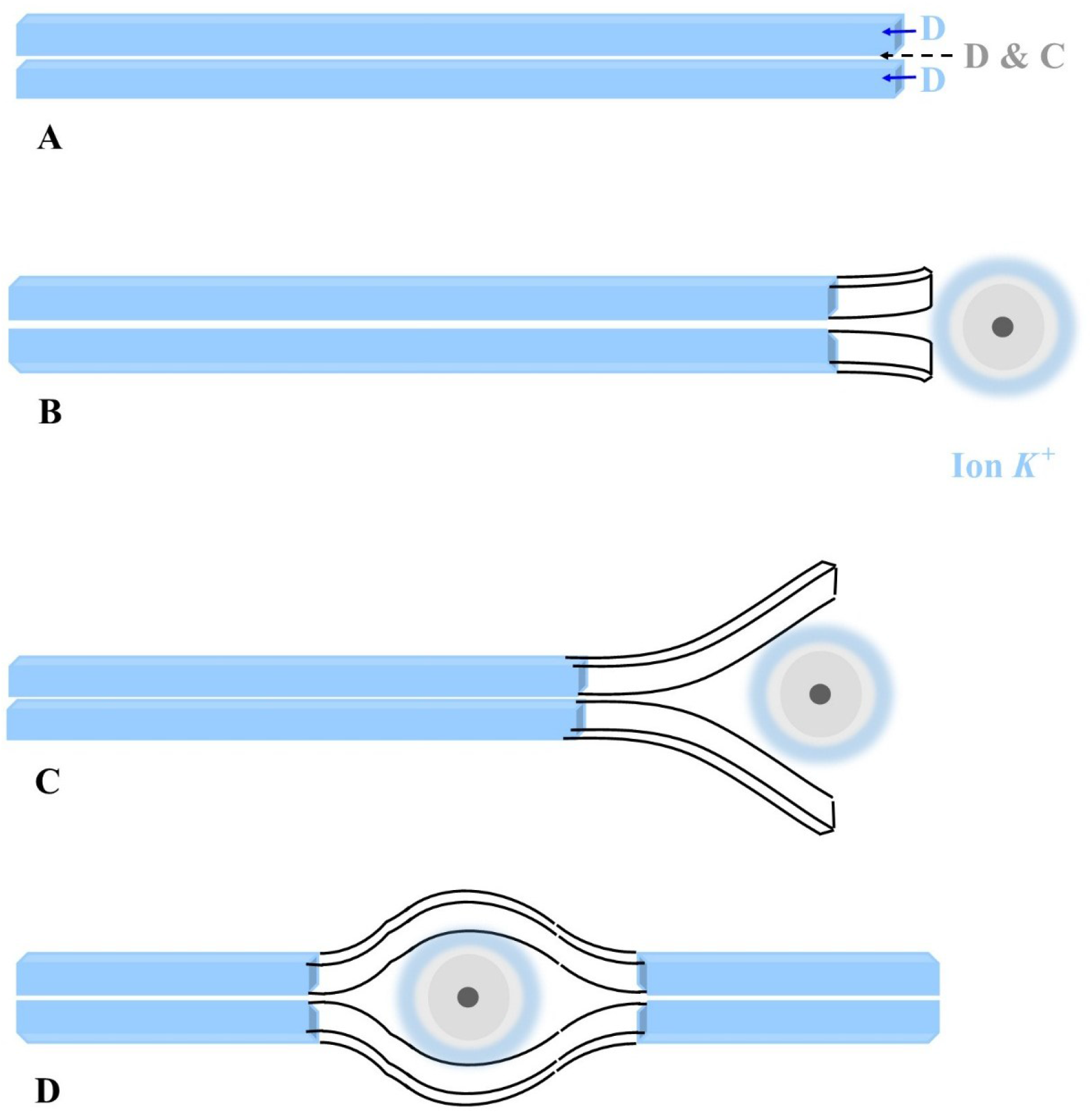
Surface waves and *K*^*+*^ tunneling through the collapsing void. The ion *K*^*+*^ is in pastel blue-grey (quantum atomic model). A: Two single myelin layers flatly pulled together with virtual void in between. D, is for dielectric, and D & C, are for dielectric and charge-trap. The virtual space, intraperiod line, *V*_*IMI*_, behaves as a dielectric, at full compression, and then as a charge-trap after capturing the electron *K*^*+*^. B and C: the ion *K*^*+*^ presents itself, the two myelin layers start to open up. D: the ion *K*^*+*^ is fully captured in between two myelin layers and remains trapped.

**Fig 14.**
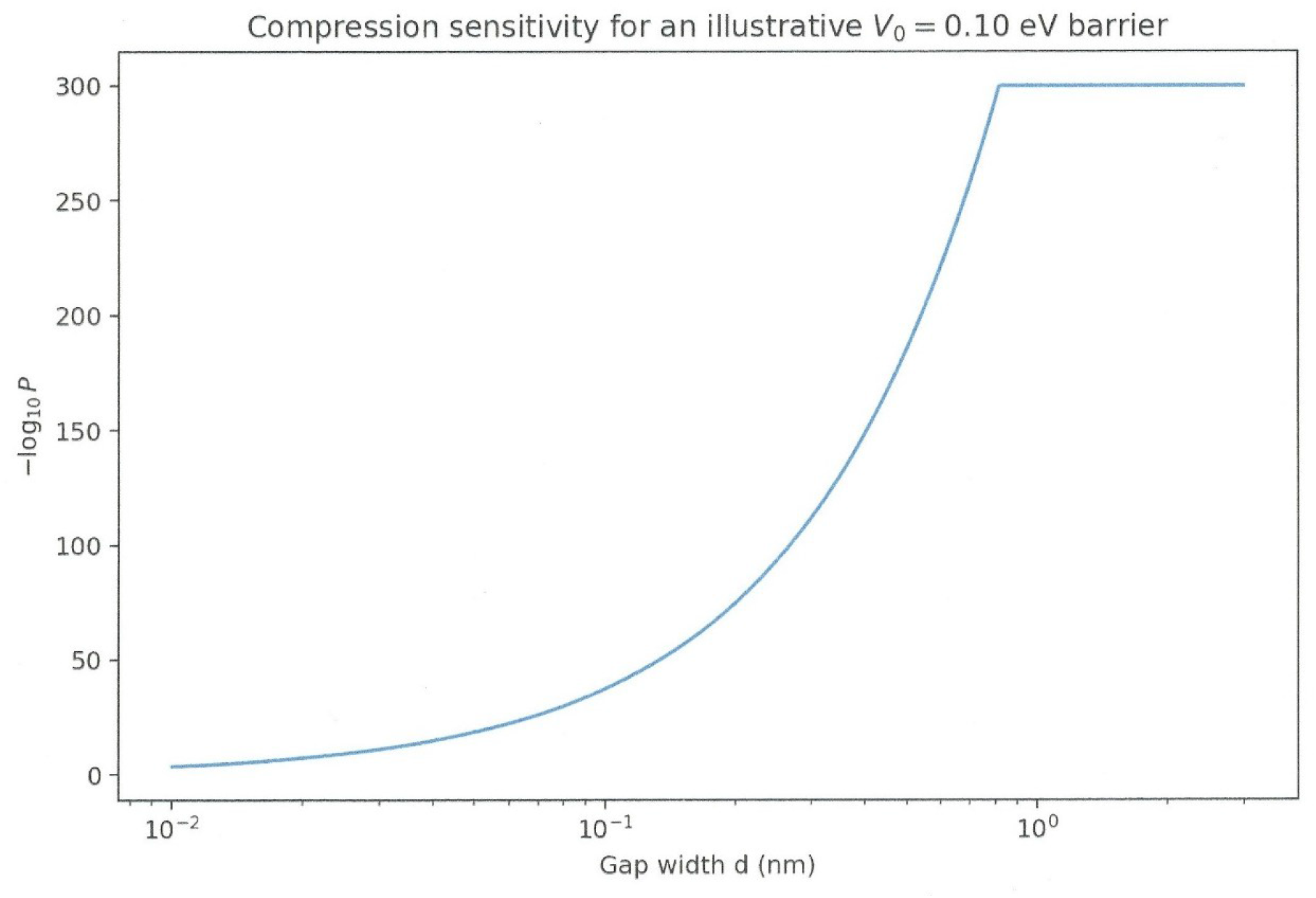
Compression sensitivity vs. initial gap. When the initial gap is almost closure in the ångström range, the quantum probability rises to 0.1-1. *d* ⟶ 0.001 nm, *P* ⟶ 1. This may occur when full compaction of myelin occurs. When myelin is not compacting and becomes loose such as neurodegenerative diseases (Alzheimer’s), then memory is dramatically affected.

**Fig 15.**
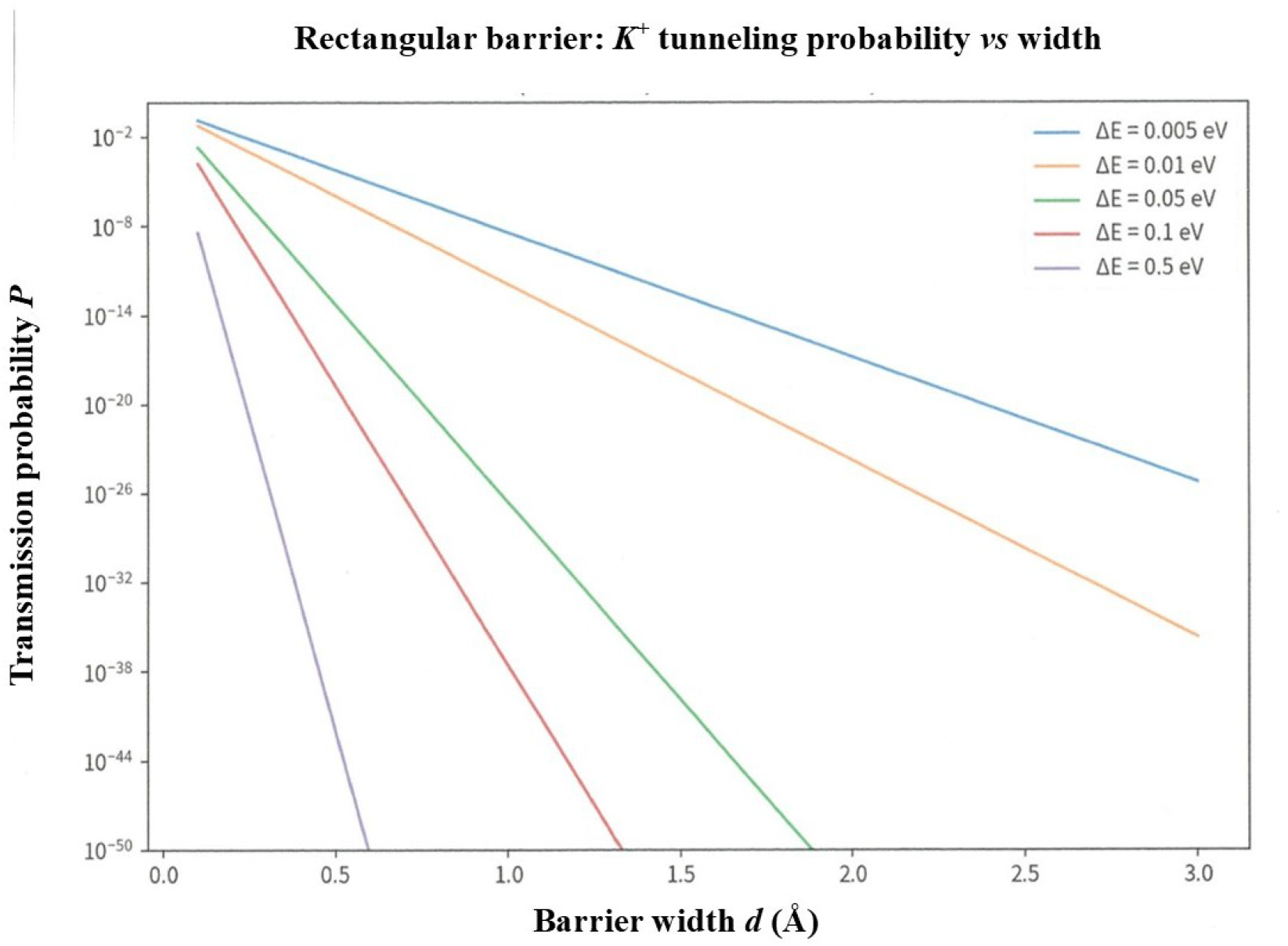
Tunneling probability through a rectangular barrier with no decoherence..

**Fig 16.**
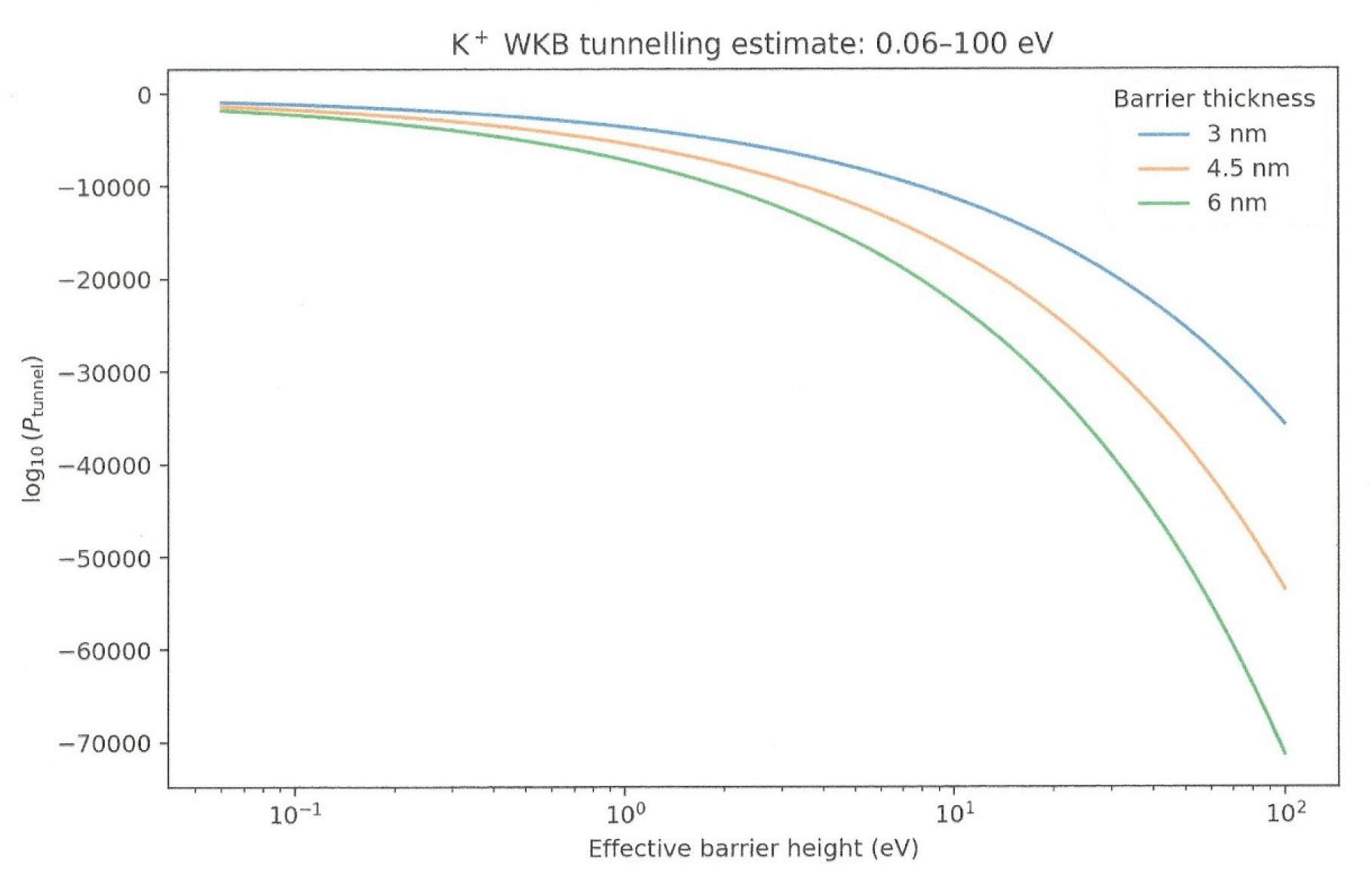
Probability of quantum tunneling *vs*. barrier height. WKB tunneling estimate.

**Fig 17.**
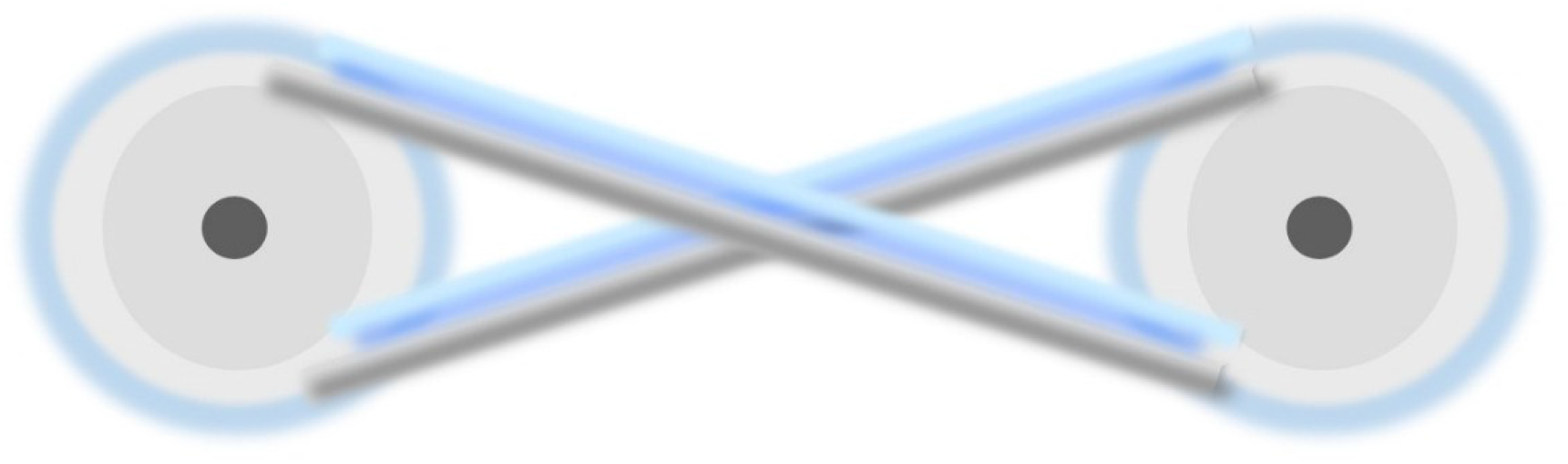
Resonant tunneling between two ions *K*^*+*^ through the intraperiod space compartment volume, *V*_*IMI*_, could behave as a quantum well to sandwich the ions *K*^*+*^. Both each extremity, “right”, and “left”, are potential barriers and sandwich the ions *K*^*+*^ which enter in resonance. The ion *K*^*+*^ is in pastel blue-grey (quantum atomic model).

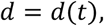

and consequently,

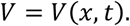

That makes this a time-dependent tunneling problem rather than a static rectangular barrier calculation we considered previously.

#### ii. *K*^*+*^ tunneling through the collapsing void

In principle, it is possible. A vacuum/void region can constitute a quantum-mechanical barrier. However, there is a major caveat. A bare vacuum cap is not automatically a high potential barrier for a *K*^*+*^ ion. The relevant quantity is the actual potential-energy landscape of the *K*^*+*^ center of mass, including:

- Electrostatic potential from the local field,
- Polarization/image-charge effects at the two dielectric interfaces,
- Energy of *K*^*+*^ relative to the two insulating surfaces,
- Changing separation of *d*(*t*),
- Ion *K*^*+*^ incident kinetic/quantum energy.

Hence, we cannot yet assign a numerical barrier such as 0.1 eV to our system. The numerical calculation below is an illustrative sensitivity analysis showing what compression does once barrier *V*_0_ is specified.

#### iii. Myelin layers’ compression

The AP is accompanied by mechanical axonal waves creating fast and temporary changes on radius, pressure, optical properties, release-subsequent absorption of small amounts of heat, and axon shortening^24^. These surface waves are driven by the travelling wave of the AP-electromagnetic depolarization, modifying compressive electrostatic forces across the membrane^24^. A periaxonal nanocircuit, relying on myelin layers compaction was recently identified^25^.

Suppose the barrier can approximately be represented by

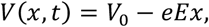

over

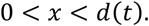

For a *K*^*+*^ ion, *m*_*K*_: is the *K*^*+*^ mass, *m*_*k*_ ≃ 39.10 u ≃ 6.49 × 10^−26^kg.

The local electric field is

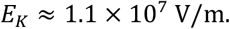

The field contribution to the potential energy is

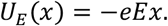

Therefore, the barrier becomes progressively thinner as

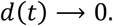

A picometer-scale compression of two myelin layers to almost perfect void, can matter enormously for *K*^*+*^ tunneling.0.

#### iv. WKB tunneling probability

For one-dimensional barrier, the WKB, tunneling probability is approximately

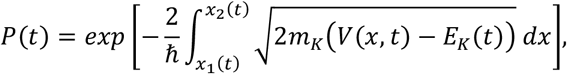

where *x*_1_(*t*), and *x*_2_(*t*) are classical turning points. This equation immediately shows why compression can have enormous effect. The exponent contains the integral over the barrier width. For one-dimensional barrier

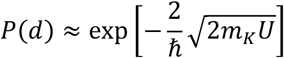

As

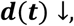

Also

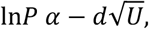

and consequently

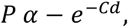

for constant *U*.

the action

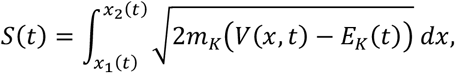

decreases.

Since

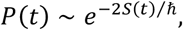

Even a relatively modest reduction in *S* can produce an enormous increase in *P*. Hereby, tunneling could increase exponentially during compression.

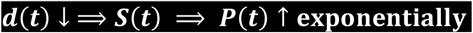

provided the ion *K*^*+*^ remains in the tunneling regime.

#### vi. Tilted, time-dependent and collapsing barrier

As aforementioned, the electric field, *E*_*K*_ is intense,

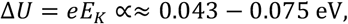

Therefore, the electric field tilts the barrier substantially, but it does not make the barrier transparent and disappear. For a tilted barrier

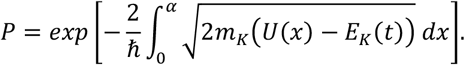

Rather than using your previous rectangular barrier, we may use

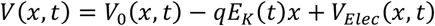

Where,

- *V*_0_(*t*): instantaneous effective barrier,
- *q*: +*e* for *K*^+^,
- *E*_*K*_(*t*) ≃ 10^7^ V/m,
- *d*(*t*): instantaneous gap,
- *V*_*Elec*_ (*x, t*): represents additional 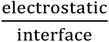 effects.

For the first sensitivity calculation, we can temporarily set

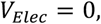

and explicitly identify this as an effective barrier model, rather than pretending that experimentally known. The two classical turning points satisfy

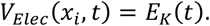

The WKB action is therefore,

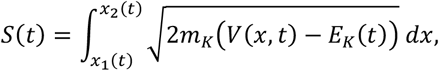

A *E*_*K*_(*t*) ≃ 10^7^ V/m field does strongly tilted the barrier.

#### vii. Sensitivity analysis

The barrier height, 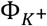, is an unknown parameter, and we calculated log_10_*P* values. The calculation predicts an enormous transition in sensitivity as *d*(*t*) approaches the sub-nanometer regime.

**Table 2.** Calculations of log_10_*P*, as a function of 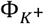 in meV, and *d*(*t*) in nm.

| $\Phi_{K^+}/d(t)$ | 3 nm | 1 nm | 0.5 nm | 0.1 nm | 0.01 nm |
| --- | --- | --- | --- | --- | --- |
| 1 meV | -2.50 | -2.50 | -2.50 | -2.50 | -0.37 |
| 5 meV | -28.0 | -28.0 | -28.0 | -7.97 | -0.84 |
| 10 meV | -79.2 | -79.2 | -51.2 | -11.6 | -1.19 |
| 50 meV | -661 | -252 | -129 | -26.4 | -2.66 |
| 100 meV | -1038 | -366 | -185 | -37.5 | -3.76 |
| 500 meV | -2482 | -836 | -419 | -84.0 | -8.40 |

At maximum compression, the gap, *i*.*e*. intermyelinic intraperiod compartment volume *V*_*IMI*,_ is closing to sub-nanometer regime, *d*(*t*) ≈ 0.01 nm, at atomic angström scale, quantum tunneling transmission may occur, ion *K*^*+*^ is forcing entry and the two deformable myelin layers are bending to let it through. The probability *P* of quantum tunneling becomes

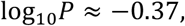

which is approximately

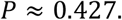

This is a stretch requiring highly restricted parameter threshold. At 5 meV, *P* ≈ 0.1415, At 10 meV, *P* ≈ 0.0646. This is not a claim of a transparent barrier height. However, in each paranode/juxtaparanod-system, there are *∼* 10^7^ − 10^8^ ions ion *K*^*+*^ available for quantum tunneling. The brain contains approximately 87 billion neurons (SD = 8), the total number of nodes would approximately be ranging from *N* ≈ 87 × 10^9^ × 10^*x*^, with x varying, *e*.*g*. for one hypothetical axon 10 mm long, *x* ≈ 10 − 50 nodes, and *N* ≈ 87 × 10^10^ − 435 × 10^10^ total nodes. Each node is shuffling *∼* 10^7^ − 10^8^ ions *K*^*+*^. Therefore, the total amount of ions *K*^*+*^ that can be shuffled at instant in the brain is 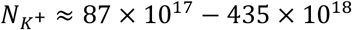. Hereby, there is a mixing of ultra-big numbers with ultra-small values. A mitigation or modulation may occur.

#### viii. Interesting possibilities

These options are different from the previous static tunneling. The steps are

AP/Node **1** ⇒ Gap**1** opens ⇒ *K*^+^approaches ⇒ *K*^+^enters transient nanoscale region ⇒ the **void *d***(***t***) decreases

⇒ the *K*^+^wavefunction encounters a **time** − **department** barrier ⇒ gap begins closing

⇒ probability of transmission can increase extremely rapidly as ***d***(***t***) decreases

⇒ K^+^crosses forcing entry between two myelin layers having very low resistance to bending and highly deformal

⇒ gap closes ⇒ leaving *K*^+^in a charge − trap region ⇔ quasi − bound resonance ⟸ reflection

⟸ Gap**2**

There are actually two fascinating physical options for the transfer of *K*^*+*^, prior to the capture or trapping after crossing. Note that the tunneling probability is extraordinary sensitive to the actual time-dependent potential barrier, not merely to the nominal gap width. However, an empty vacuum gap is not automatically a low-energy barrier for ion *K*^*+*^ ion. For a massive ion, the decisive quantity is the *K*^*+*^ potential energy profile *V*(*x, t*), including the myelin interfaces, electric field, and the ion’s initial energy. A gap of 0.1 − 3 nm does not by itself tunneling. Myelin layers are highly deformable under force and can easily bend to allow ion *K*^*+*^ to passing through. Myelin wrapping is flexible^26^. At a molecular scale, myelin’s lipid layers have a very low resistance to bending, *aka* “bending rigidity”. This elasticity allows individual layers to smoothly accommodate changes in local curvatures. Because the tunneling probability depends exponentially on the semi classical action, progressive reduction of the instantaneous forbidden region can produce extremely rapid, approximately exponential increase in transmission probability during gap closure.

#### ix. Quantum tunneling under compression

If

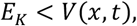

while the barrier remains finite, then the wave function can penetrate the barrier,

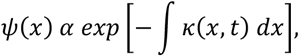

with

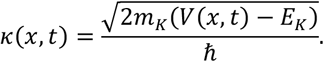

Then compression can dramatically increase the transmission probability.

#### x. Capture after crossing

If the *K*^*+*^ emerge on the opposite side, and the two insulating myelin layers subsequently close around it, the ion *K*^*+*^ could potentially become spatially trapped. However, the trapping must be specified, to enable metering, sorting, organizing the data for retrieving the appropriate and relevant data at a point of time. This trapping could be,

- Electrostatic trapping,
- Polarization/image charge trapping,
- A potential minimum,
- Steric confinement,
- Or some combination.

#### xi. Compression inference

If

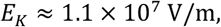

and

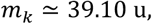

this mass is a major quantitative issue. For an electron, tunneling over nanometer distances can become readily important. The ratio

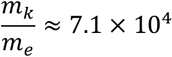

The WKB contains 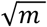.

Therefore, the mass penalty is approximately

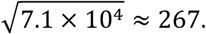

Hereby, an otherwise similar *K*^*+*^ barrier is vastly more suppressive than an electron barrier. That does not make tunneling mathematically impossible. It makes tunneling occurring only under highly restricted parameter threshold; the barrier has to become exceptionally narrow and/or extremely low. However, should this occur, under the voltage-controlled fine tuning on the *K*^*+*^ filtering, the metering and organization of the data/*K*^*+*^ trapping are maximized.

We examined a simple linearity tilted barrier as an illustrative model.

For

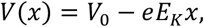

the WKB integral can be evaluated analytically,

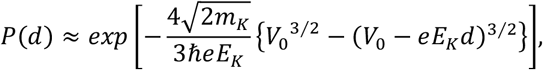

provided that *V*_0_ > *eE*_*K*_*d*. This equation is particularly useful for the compression hypothesis because *d* occurs explicitly in the exponential.

**Table 3.**
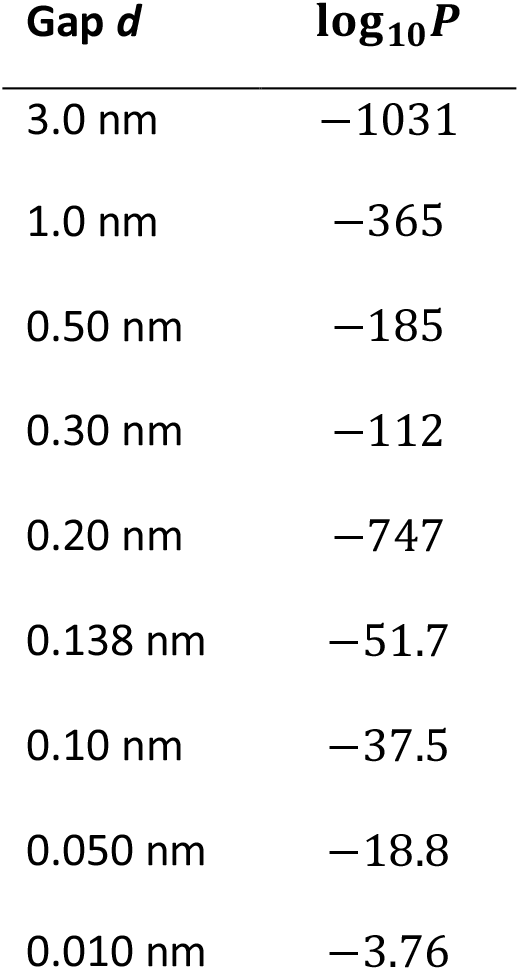
Shows that the more compaction, and suppression of the void, the more probability exists for quantum tunneling. This is counterintuitive.

#### xii. Numerical example

We evaluated the model for illustrative barrier heights,

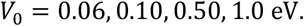

These are not measurements of the structure while it is compressed. They are sensitivity analysis showing how strongly the results depends on the unknown values for the actual barrier.

Here are the results for *V*_0_ = 0.10 eV,

In this illustrative model, reducing the gap from 1 nm to 0.1 nm changes the predicted probability by roughly

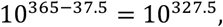

in relative scale. That is an enormous exponential sensitivity. It also demonstrates something important, and intuitively paradoxical:

**For an ion *K***^***+***^, **quantum tunneling through a nanometre-scale vacuum gap is extraordinarily suppressed unless the effective barrier is extremely low and/or the gap becomes extraordinary small**.

Therefore, the compression mechanism is potentially powerful, but the actual *V*(*x, t*) is absolutely critical.

#### xiii. The ion *K*^*+*^ radius factor

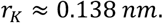

Although this is a geometrical value, we must be careful using it as is as a tunneling width. In quantum mechanics, the relevant coordinate for WKB is the center-of-mass coordinate of the ion, not simply its ionic radius. Hereby, it does not mean that *K*^*+*^ automatically has a *d* = 0.138 *nm* tunneling barrier. However, it gives us an important physical scale for determining when the two surfaces become extremely close to the ion *K*^*+*^.

#### xiv. The exponential acceleration

Compression reduces the WKB action, producing an exponential increase in transmission probability.

The distinction matters because the probability changes exponentially. The actual number of transmitted ions *K*^*+*^ per unit time depends additionally on the rate at which ions *K*^*+*^ encounter the barrier.

#### xv. How many ions could cross?

If

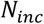

Ions encounter the barrier during a particular AP event, then the expected number transmitted is

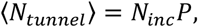

For independent identical AP events. More generally, if the arrival rate is

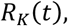

Then

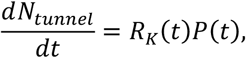

and therefore,

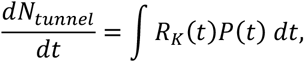

this is an important equation, because *P*(*t*) would rise enormously during compression.

#### xvi. Numerical example for an AP event

Suppose, purely illustratively,

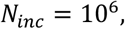

then for a small gap,

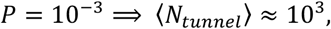

whereas for a larger gap,

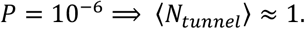

Hereby, even if the effect is small, the number of ions *K*^*+*^ that can cross,

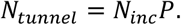

We can only obtain the absolute number *N*_*tunnel*_ until *N*_*inc*_ or equivalently the ions *K*^*+*^ arrival is known.

#### xvii. The environmental assumptions

They will be discussed one by one in detail in the Discussion section. We have deliberately specified a near-ideal topological insulated myelin system. New experiments and new findings shed light on the unique myelin system. However, this system is at the edge of the threshold quantum coherence-decoherence during “normal” homeostasis. Because myelin alterations are the hallmark of neurodegenerative diseases (Alzheimer’s, multiple sclerosis, etc.), the system is losing quantum coherence, settles in permanent decoherence and is no longer able to achieve near-ideal topological insulation. For the system to work, it requires a perfect myelin structure and myelin structural adhesion, *e*.*g*. myelin basic protein (MBP). Those assumptions for a coherent myelin system are as follows.

- No water
- No residual gas,
- No thermal fluctuations,
- No phonons
- No excitations in the surrounding material,
- No fluctuating electromagnetic fields,
- No electrode/insulator degrees of freedom,
- No spontaneous electromagnetic emission,
- No charge-position fluctuations,
- No external environmental coupling
- No degrees of freedom associated with the mechanism driving the ions.
- The two ensembles of S_1_ and S_2_ of *Na+*, 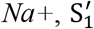 and 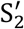 of *K*^*+*^, at each extremity of the cylinder, distance L between them, are separated at all times.

If those homeostatic conditions can be achieved, they make the theoretical problem much cleaner. The system approaches something like

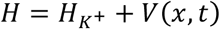

with the time dependence primarily coming from the mechanically changing geometry. The Schrödinger equation becomes

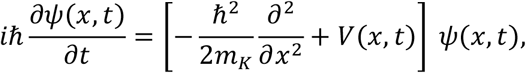

which a more rigorous expression for this option.

#### xviii. The moving barrier

Because the myelin layers are compressing,

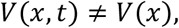

the time-dependent numerical Schrödinger calculation is preferred over the static WKB approximation. For example, if

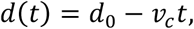

where *v*_*c*_ is the compression velocity, then

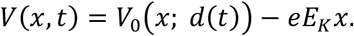

The transmission probability becomes

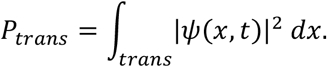

This would let us explicitly simulate,

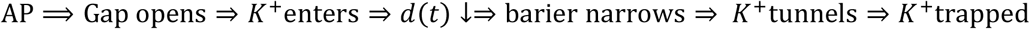

This is a much stronger model than simply assigning a fixed *d*.

#### xix. Resonant tunneling

The local magnetic field value in the intraperiod myelin insulated system is not yet available. Measurements of the extra cranium overall values does not bring the useful information needed. The intermediate region cannot be assumed in resonant-tunneling state unless the calculated potential actually supports a quasi-bound eigenstate or a resonance pole. The system 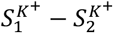 seems to work this way. There is a tunneling barrier at the entry and the mirrored structure at the other node, and the in between intraperiod space compartment volume, *V*_*IMI*_, could behave as a quantum well to sandwich the ions *K*^*+*^. Both each extremity, “right”, and “left”, are potential barriers where ions *K*^*+*^ could reflect. Although it seems logically possible, we do not yet have evidence.

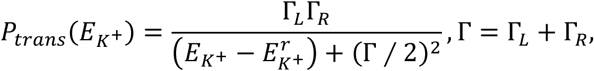

where Γ_*L*_ and Γ_*R*_ are energy widths, not lengths and not probabilities. They describe how rapidly the intermediate quasi bound state leaks out.

- Γ_*L*_: coupling/leakage form the quasi bound state toward the “left” S_1_ side (Node 1).
- Γ_*R*_: coupling/leakage toward the “right” S_2_ side of the cylinder (Node 2).
- Γ = Γ_*L*_ + Γ_*R*:_ total resonance width.
- 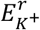: resonance energy.
- 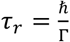: associate lifetime of the resonance.

Thus a narrow resonance has small Γ, long lifetime and a very narrow energy window. A broad resonance has larger Γ, short lifetime and a broad energy window. For a symmetric coupling,

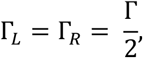

And the ideal one-dimensional Breit-Wigner peak reaches

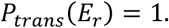

This is important, because resonant tunneling can turn exponentially a tiny off resonance transmission into an order unity transmission, but only if a genuine intermediate state exists and the incoming energy is sufficiently close to (*E*_*r*_).

Note that, *L*, distance between two nodes is not the tunneling distance. Also,

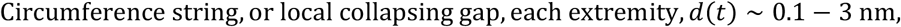

and,

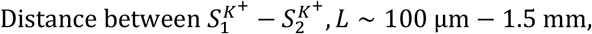

and

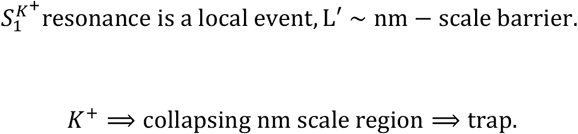

The 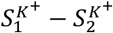 interaction is instead a long range field mediated problem,

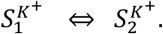

Hereby, there is no reason to insert long range L into the tunneling exponential for the 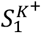 event. That would make it astronomically smaller.

At resonance,

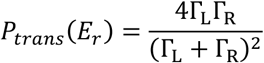

And therefore,

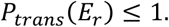

The off-resonance value at detuning is

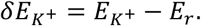

The enhancement factor is

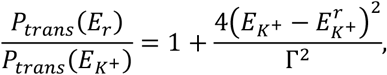

For a symmetrical ideal case.

Therefore if

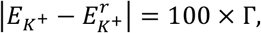

Then,

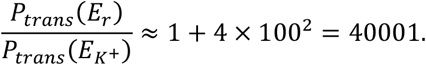

If,

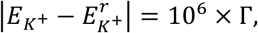

Then the ratio is about

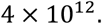

The resonant-tunneling hypothesis is legitimately testable and the resonant enhancement can be enormous by many orders magnitude. However, this enhancement requires

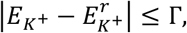

During the relevant par =t of the compression trajectory.

#### xx. picosecond, and instantaneous on-off

Since

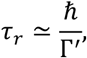

For a resonance lasting 1as,

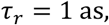

Would imply

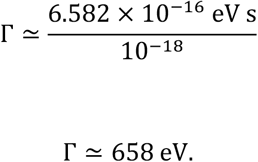

For 2 as,

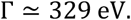

Those are extremely broad energy widths. By contrast, a resonance width

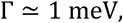

has

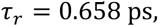

and,

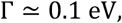

gives

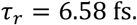

A 1 − 2 as, triggering time does not automatically induce a 1 − 2 as, resonance time. Those are two different physical quantities. The “all-or-none” phenomenon really represents a phase locking/correlation between 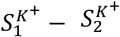. The intermediate *K*^*+*^ quasi-bound state itself does not last 1 − 2 as.

For the brain myelin system, it is useful to separate the Hamiltonian (*K*^*+*^ ions) into three contributions,

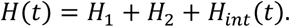

The density matrix (*K*^*+*^ ions) is

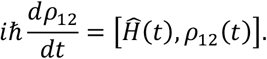

For an electric field-only model, 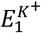, at Node 1 (left), and 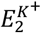, at Node 2 (right), where the value of local intraperiod-intermyelinic magnetic field not yet available, a minimal two-mode collective model is

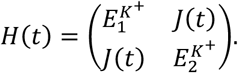

Here *J*(*t*) is the effective 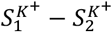.

This gives,

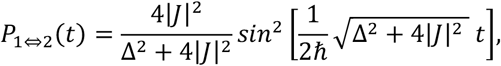

For a simple constant coupling two-mode model, where

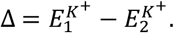

This is describing the coherent coupling between two collective modes, albeit this is not a tunneling formula. Note the following essential distinction,

**phase synchronization # resonant tunneling**

and

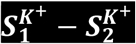 **coupling # local *K***^***+***^ **quasi-bound state**

**1 2**

Does a quasi-bound state exist? This proposed hypothesis is genuinely testable, albeit some parameters are missing.

#### xxi. The phenomenon is experimentally testable

The entire phenomenon would be measurable and testable such as the number of ions *K*^*+*^ transferring. Quantum tunneling of *K*^*+*^ through a transient, collapsing nanoscale vacuum gap is allowed by the Schrödinger equation and can be strongly enhanced by the time-dependence and by resonance. However, the exact numerical calculation could only be obtain if we know some parameters that do not have yet been measured, such as *v* _*c*_, *d*(*t*), *d*_0_, *V*(*x, t*), 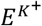, *L*, Γ _*L*_/Γ_*R*_. Quantum tunneling of ions *K*^*+*^ showed an extreme sensitivity to those parameters. The overall sensitivity analysis shows how the quantum tunneling probability is enhanced exponentially, by many orders of magnitudes when those parameters are within a at highly restricted parameter threshold range. The effect sits on a knife-edge of parameter space. Realistic numbers of for the final gap, magnetic field, and barrier height are still missing. Therefore, only the functional dependence and the enormous sensitivity in the razor-edge space of parameter can be stated with certainty. This geometry is experimentally accessible and would constitute a striking demonstration of macroscopic quantum dynamics of heavy ions if this transmission window can be realized.

**Table 4.** Sensitivity analysis with influencing parameters.

| parameter | direction increasing P | sensitivity |
| --- | --- | --- |
| **final Gap $d_0$** | minimize (target $\leq 0.15$ nm) | exponential $\uparrow\uparrow\uparrow$ |
| **barrier height $V(x, t)$** | minimize (via field bending) | exponential $\uparrow\uparrow\uparrow$ |
| **local field $E^{K^+}$** | maximize (stronger tilt) | strong $\uparrow\uparrow$ |
| **compression trajectory $d(t)$** | faster final collapse while wave packet is present | high ↑ |
| **resonance condition** | align incident energy with quasi-bound level | Can give orders-of-magnitude boosts $\uparrow\uparrow\uparrow\uparrow\uparrow$ |
| **ion kinetic energy** | optimize relative to instantaneous, $V(x, t)$ | moderate |

Those parameters are definitely measurable and will probably be in the near future. The equations of quantum mechanics are exact, they are exceptionally accurate models of reality that have passed every single experimental test for over a century. A direct measurement of quantum mechanics is difficult although possible. In the case of the brain, the physiology and definition of boundaries must be exact and complete. A single missing or inexact element of information of anatomy, physiology, biochemistry, cytology, histology, and nano leads to an inaccurate model of the myelin system and the quantum mechanics equations are wrongly apply to false boundaries.

Let’s pursue our evaluation, and perform a sensitivity analysis, playing with parameters we do not yet know. For example, measure

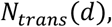

while systematically varying *d*. A tunneling-dominated regime should approximately exhibit

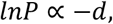

for a simple rectangular barrier, or the corresponding nonlinear relationship for a field-tilted barrier. For this dynamic hypothesis, an even more interesting measurement would be

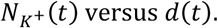

The *K*^*+*^ transmission increases dramatically as the two layers approach one another.

Under the vacuum, low-bending rigidity, and near-ideal topological insulation, of the intraperiod myelin system, *V*_*IMI*_, quantum tunneling of ion *K*^*+*^, through the transient collapsing gap is allowed by quantum mechanics and can be dramatically enhanced by the time-dependence of the barrier, and further by resonance. Quantum tunneling is in principle possible, but under normal idealizations the transmission probability is expected to be extremely small, mainly due to the size of the heavy ion *K*^*+*^, radius *∼* 0.138 nm, and mass *∼* 7 × 10^4^ times the electron mass. The transmission is more suppressed than for the electron. Except for highly restricted parameter threshold conditions, the transmission probability can increase exponentially and massively. Those threshold conditions are: 1. narrowest gap *d*(*t*) ≲ 0.1 − 0.3 nm; 2. A tilted barrier; 3. A compression of gap; 4. Resonant tunneling; 5. Enormous number of paranodes, juxtaparanodes involving huge number of ions *K*^*+*^, 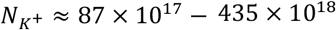. However, only a much lesser number of ions *K*^*+*^, would be recruited during one impulse. The repeating impulses for one information to be retained and store, may have a cumulative effect of synaptic strengthening which has been widely reported, *aka* classical Hebbian effec^t1,27-31.^ Under the most optimistic combination of the above parameters, a single-ion transmission probability is in the range of 10^−12^ − 0.4, or even higher if a strong resonance is hit. If this occurs, the filtering, of the ions *K*^*+*^ capture into the charge-trap *V*_*IMI*_, this hypothesis becomes conceivable during a brief temporal window. It would allow the fine metering voltage-controlled each impulse allowing the sorting, organizing and further retrieving of the data stored in the *V*_*IMI*_ well. The absolute number that could tunnel is therefore enormous, in principle. In practice, the mutual Coulomb repulsion, the finite duration of the open window, and the requirement that the gap closes after the ion *K*^*+*^ has crossed limits the yield. Once the layers come into contact (near contact), a sufficiently deep local potential minimum can trap the ions that have passed; the depth of the trap relative to residual kinetic energy added to the Coulomb energy determines the long-term retention.

#### xxii. Experimental distinction

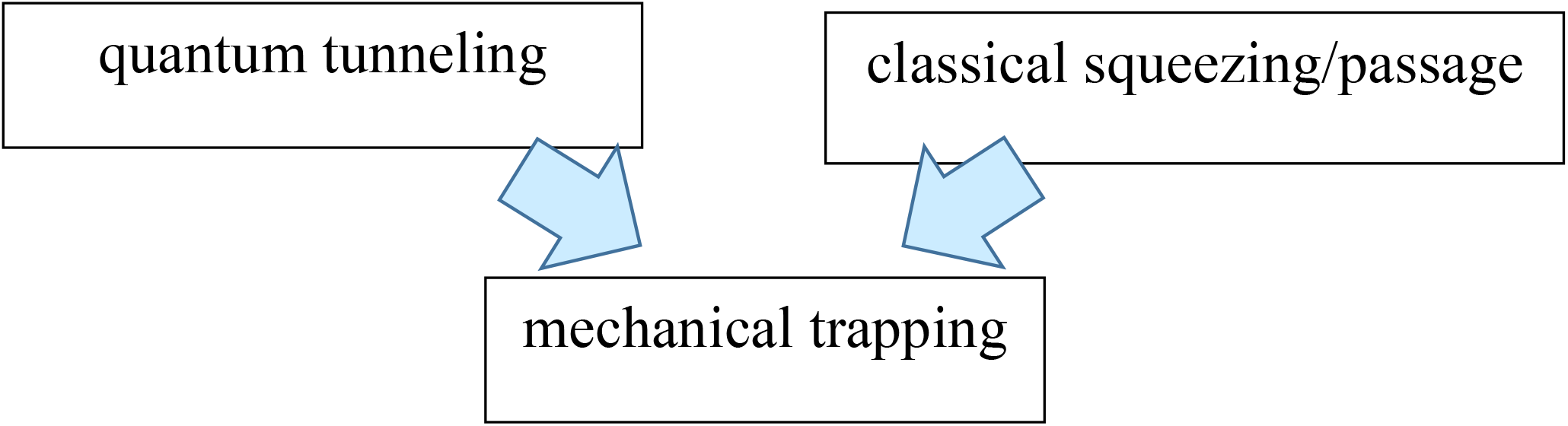

Another interesting sensitivity analysis would hence measure both the gap trajectory. Conceptually

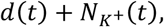

is much more informative than measuring *K*^*+*^ accumulation alone.

### C-III. Classical passage

If the gap is sufficiently open, and ions *K*^*+*^ have enough energy, classical physical laws apply,

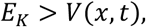

Then the ion *K*^*+*^ simply crosses classically. Note that this is not quantum tunneling. Although without the compression-induced exponential transmission probability, there is no exiting force such as quantum tunneling enabling the forced entry between the two myelin layers. We are unable to assess the nature of the driving potential, voltage or force that would be required to open the gap of the intraperiod myelin system, *V*_*IMI*_, neither would we to suggest a force to cross the enormous barrier height without quantum tunneling and the other parameters decreasing the barrier height (compression, resonance, etc.). The second caveat with classical passage is the metering of data. Because quantum tunneling is organizing individual charges into separate bins like sorting mail, quantum tunneling “sorts” data by metering the exact quantity of charges (electrons) allowed to pass through a quantum barrier^18-20^. This capacity is likely to be lost should classical physics applies. Classical physics via mechanical motion of myelin layers or other physical principle cannot create a system of metering equivalent to a quantum barrier and the precise AP voltage.

## D. Discussion

### D-I. A potential candidate for near-ideal topological insulation

#### i. Exclusion of implausible models for physiological memory storage and engram inference

We analyzed and inventoried the significant progress that has been made in elucidating the cellular machineries underlying memory formation^1^. This inclusive and thorough analysis of memory formation are the pillars providing deeper insight to further elaborate on physics and physiology^1^. Indeed, the precise physiological, molecular, and physical (quantum) substrates of memory storage, and location, remain an open question^29-33^. *It does not exist a single specific protein nor single specific gene encoding for a single specific information-memory bit*. The principle of exclusion (Michelson–Morley) is guiding to cancel implausible biological models underlying the storage for long-term potentiation of memory. Hereby, we are left with a finite scope of physiological and physical paths^1^.

Indeed, DNA is a reliable and stable structure to store data on the long term, resisting time even millennia. Using the CRISPR-Cas system, it is possible to write information encoding pixel values in the genome of living cells, such as digital short films^34^. However, the timeframe required to generate lasting hard mutations in oligonucleotides structures across generations is too long to explain the learning process in humans. Holocaust exposure induced intergenerational epigenetic changes on *FKBP5* methylation, without any hard alteration of the underlying DNA codes^35^. Epigenomics seemed to have programming next generations’ fear and survival response to earlier PTSD. However, in the fast learning timeframe of childhood and later, DNA and epigenomic acquisitions do not seem to play such reliable, massive and rapid potential storage. This might have molded a favorable evolution of brain structure to innate functionality on the long term. The human brain has several specialized processing centers filtering sensorial data, analyzing, sorting information. Memory consolidation implicates a widely distributed network beyond the hippocampus^33^. Perception, motion or navigation in a three-dimensional space depend on spatial memory and require permanent visual tracking of the direction and distance from reference points, or GPS-like landmarks, and their integration by hippocampal and entorhinal circuits supporting grid-cell-based spatial mapping^36,37.^Time boundary cells, specialized neurons in the entorhinal cortex, hippocampus, and lateral prefrontal cortex (LPFC) are internal temporal organizers of memories segmenting working-memory and navigation epochs necessary for recalling goal-directed paths^38^.

Those brain AI-like processing centers may sort, organize, and triage information which is later stored in the intraperiod-myelin system we are describing. White matter constitutes approximately 40-60% of the total brain volume which is readily available for transmission and potentially storage. The blood-brain barrier, BBB, isolates neurons and white matter tracts from the circulation, compared to other body cells. BBB with its tight sealing provides a first frontline protection of chemical, functional distance, and strict environmental control. Padding by superposed hydrophobic myelin wrappings may provide further insulation, such as electromagnetic, thermal, even creating virtual void, and expelling water.

#### ii. Equipment inventory of the brain machinery for the engram

Let’s pursue the inventory of the physiological, biochemical, and physical equipment of the brain started in previous sections. A single oligodendrocyte can wrap and insulate up to 40-50 axon segments. The structural integrity of myelin layers is necessary for learning and memory^39^. Myelin layers are compacted to a thickness proportional to the axon diameter^40^. A single axon can have from 10 to 160 and up to 250 or 300 layers for high-speed transmission with an average range from 40 to 100. The thickness of a single myelin layer ranges from 3.9 to 20 nm. The dimension scale is in nanometers, approximately the exact dimensions of the charge trap, the tunnel and dielectric also equipping current AI microchips, type NVDIA, Intel, Tesla AI-5, etc., going thinner and smaller with a trade-off of damaging the dielectric membrane. On the axonal transversal slice, Figure 6a, we see the cylindrical superposition of myelin layers. The internode length typically ranges from 50 μm to 750 μm with nodes about 1-2 μm. The internode-length is sufficient to enable rapid transmission of the electromagnetic potential to the next node. The superposition of multiple, concentric, spiraled, and highly compacted cytoplasmic membrane wraps are assembled by the oligodendria (oligodendrocytes) (Fig. 6a). This spatial packed wrapping in 3D creates a disjointed multiple assembly; nodes not facing from a layer to next (Fig. 6c). This is in fact amazingly beneficial for perfect isolation emulating the memory cells of the current AI chips, collectively consisting of a massive array of cells, stacked in hundreds of layers and organized into tens of thousands of rows and columns. Individually, enabling rapid conduction with sufficient internode length.

Let’s look at the number of Ranvier nodes. The number in a single neuron varies^41^, depending on the axon length, ranging from a few to hundreds or thousands. In the human brain, for an average of 87 billion neurons (SD = 8), the total number of nodes would approximately be ranging from N = 87 × 10^9^ × 10^x^, hereby, x = (3, 4), and N = (87 × 10^12^; …, 87 × 10^13^) ≈ (90 × 10^12^; …, 90 × 10^13^) ≈ (1 × 10^14^; …, 1 × 10^15^). This would approximately be one trillion to hundreds of trillions. Figure 6b illustrates the fused layers of oligodendrocyte-insulators wrapping the axon. The myelin is constituted of lipids, highly insulating. There are minute-masses of cytoplasm trapped between fused layers of cell membranes of oligodendrocytes. Those virtual spaces of relative “negative or lower pressure points” are important for adhesion and trapping (major dense line *vs*. intraperiod line).

Let’s look at the electrolytes involved in the action potential, AP. A physiological regulation of electrolytes controls intra- and extra-cellular milieus within a strict homeostatic range. The plasmatic *Na*^*+*^ normally ranges within 138 - 142 mmol/L, and plasmatic *K*^*+*^ within 3.5 – 5.0 mmol/L. In terms of electromagnetic potential difference, sodium and potassium are the two main ions involved in the axonal depolarization and repolarization of neurons (Sections *“Transversal transfer* of charged ions across the axon, *Longitudinal jump* of electromagnetic potential along the axon”). This homeostatic control is vital for the organs and cells. The loss of electrolyte balance will impact with severe life-threatening consequences to the heart, all organs including the brain, and all cells. This intra-, extra-cellular, and plasmatic electrolyte balance of *Na*^*+*^ and *K*^*+*^ applies beyond the blood brain barrier, encompassing neurons, oligodendria, glia, and microglia. This is critical to maintain the normal negative resting value of neuronal membrane potential, and heretofore, the occurrence of action potential.

#### iii. Potential engram mechanism

In Figure 6c, at initiation of AP, *Na*^*+*^ enter the axon, *K*^*+*^ is exiting, the amount of *K*^*+*^ that cross the cell membrane during a single action potential is extremely small relative to the total concentration of inside the cell. The molar amount is estimated approximately to 3.3 × 10^−18^ moles of *K*^*+*^, per each action potential on a single axon and negligible. The amount of *Na*^*+*^ crossing the membrane during a single action potential is also relatively negligible compared to the total intracellular concentration, allowing the cell to maintain gradients without constant, immediate correction. That is 0.8 to 1.5 picomole/cm^2^ (or 0.8 – 1.5 10^−12^ moles) of *Na*^*+*^ cytoplasmic membrane area of the neuron. All of which cannot be considered as a homeostatic disruption.

The inference, and likely the only one to store the information, and engram the LTP, is as follows. Instead of returning intra-cellular within the axon, during repolarization, is to trap *K*^*+*^ ions into the multiple superposed myelin layers in the physical process as described in Fig. 6 and Fig. 2c (charge trap).

When the action potential propagates along the axon, positive charges (*Na*^*+*^) enter the neuron. This is equivalent to an electromagnetic field creating negatively charges outside the axon toward the outward concentric multiple myelin layers, acting as charge trap zone^42^. In this case, the myelin layers behave as a dielectric barrier and the applied electric field is strong enough, the charges extend across the Ranvier nodes, creating a non-negligible probability that the negative charges will appear on the opposite side within the multiple myelin layers (charge trap)^42-45^. This process, known as quantum tunneling^43-45^, infer that ions *K*^*+*^ may remain trapped in a certain architectural organization.

An image is stored in the charge trap flash (CTF) of a microprocessor by converting its visual data into binary values (0s and 1s) and trapping electrons in specific physical locations within memory cells. Unlike traditional flash that uses conductive floating gates, CTF uses a nonconductive layer to hold electrons in place. In the superposed multiple, concentric, spiraled, and highly compacted myelin wraps specifically assembled in 3D may allow such storage of charges by the disjointed 3D spatial superposition (Fig. 6a, 6c). In this case, stored charges are positive ions, with similar effects whether charges are negative or positive charges. The electromagnetic field will be created all the same. To “write” data, following an action potential, this voltage applies to the control gates of the myelin layers producing a ionic charge injection. This causes charges to gain energy and “tunnel” through the myelin layer across Ranvier nodes, via tunneling^43-45^, and deep into the concentric myelin multilayers (Fig. 6b). This is creating an insulated trapping of *K*^*+*^ ions isolated from the system. Because myelin layers are an insulator, the injected ionic charges (*K*^*+*^) become physically stuck in “traps” within the myelin lipid (lipoprotein) structure. The *K*^*+*^ ions may not move freely, completely trapped after the AP (electromagnetic field) ceases. Mirroring a single-bit, a single-level-cell, SLC, a trapped ionic charge (ions *K*^*+*^) may represent a “1,” while an empty cell (absence of *K*^*+*^) represents a “0”. The presence of trapped electrons increases the threshold voltage of the transistor; a similar phenomenon may occur in the brain for the ions *K*^*+*^. This brain apparatus may also behave as a current multi-bit, MLC (2 bits/cell), TLC (3 bits/cell), or QLC (4 bits/cell), such as the modern 2026 3D NAND SSD, solid-state drive to stack large number of non-volatile flash memory cells (NAND). All things considered, the brain may store multiple bits per “cell”, or minute-masses of cytoplasm trapped between fused layers of cell membranes, virtual spaces of relative “negative or lower pressure points” likely important for adhesion and trapping by precisely controlling the number of *K*^*+*^ ions trapped creating different voltage levels (*e*.*g*., 8 levels for 3 bits). Hereby, the trapping of *K*^*+*^ ions would be proportional to the AP intensity. All of which would be in line with neuronal plasticity, synaptic strengthening, after repeated use and reinforcement learning^27-30,46-49.^

If the homeostatic balance of *Na+* and *K*^*+*^ cannot be maintained in some ICU patients with life-threatening cardiorespiratory and renal failures, a normal brain function cannot be sustained and coma may result. Electrolyte abnormalities and ensuing failing action potential is one of the causes of brain dysfunction. High levels of extra-cellular *K*^*+*^ concentration may be associated to a neuronal firing pattern, mixed-mode bursting, MMB^50^, which is never attained in physiological action potential depolarization. When *K*^*+*^ ions are detrapped from myelin and oligondendria wraps, they may trigger action potentials causing depolarization. Indeed, a slight increase of extra-cellular *K*^*+*^ ion reduces the concentration gradient across the neuronal making the resting membrane potential less negative, closer to the threshold making it more excitable and prone to propagate AP^51^. A slight rise of 3mmol/L sufficiently the resting membrane potential toward the threshold making it easier to trigger and sustain AP. The Neuropixels 2.0 probe^22^, or rather, the Neuropixels Ultra (NP Ultra)^23^ measuring a single-neuron, single spike resolution, with silicon probe with much smaller and denser to probe extracellular waveform may bring further insight. To support this inference, a resolution in the nano range exploring the minute-masses of cytoplasm trapped between fused layers of cell membranes, virtual spaces of relative “negative or lower pressure points” likely important for adhesion and trapping the *K*^*+*^ ions would be needed in a complex experimental design. Studies found a mechanical displacement of the axonal membrane accompanying the electromagnetic potential^24^. This mechanical displacement potentially exposes the charge trap (Figs. 6c and 7) through the Ranvier nodes across the myelin-dielectric via a tunneling process. The potential energy is stored in elastic properties of the neuronal membrane and cytoskeleton while kinetic energy is carried by the axoplasmic fluid. These surface waves are driven by the travelling wave of the AP-electromagnetic depolarization, modifying compressive electrostatic forces across the membrane^24^. This potential difference is also propagating mechanical displacements, which the authors called “action waves”^23^. We also infer that this geometrical deformation may facilitate the tunneling and also that *K*^*+*^ ions become trapped in minute-masses of cytoplasm, joint surfaces forcing apart (tribonucleation), between fused layers of cell membranes, exposing virtual spaces of relative “negative or lower pressure points” to let the *K*^*+*^ ions enter and being trapped, then the “action wave” clips them in. Sequestering those *K*^*+*^ ions may be indefinite, until a new sufficiently AP stimulus above threshold (*e*.*g*., a memory recall) reopens those “dormant myelin-memory cells”, further detrapping *K*^*+*^ ions, generating intense AP.

### D-II. Quantum coherence preservation in the intraperiod-myelin system

Based on recent data, let’s review and weigh one by one *potential channels of decoherence*.

- *No water and no residual gas*. The biological “vacuum-sealing” creates a tightly wrapped, highly efficient insulation around brain nerves’ fibers. The forces the of the lipid hydrophobic myelin layers stick flatly and tightly against one another like a perfectly zipped jacket^2,3.^The oligodendrocytes wrap their membranes around the axon, and do not leave it loose (neurodegenerative pathologies, *e*.*g*. Alzheimer’s or demyelinating conditions, *e*.*g*. multiple sclerosis). They actively squeeze out water, extra fluid, gas, and cytoplasm between the layers. Molecular zippers (MBP), microscopic zipper or magnet neutralize negative charges on the membranes, pulling adjacent layers flatly together into “compact myelin”^2,3.^
- *No thermal fluctuations, temperature changes* in nerves were unmeasurably small, *i*.*e*. smaller than 40 μK ≈ 0.00004°C, totally reversible at refractory phase. This was at the sensitivity limit of the experiment^52-54^. While heat generally destroys quantum coherence, recent advancements in quantum physics and biology demonstrate that coherence can be preserved or dynamically maintained even in warm environments, and at or near physiological temperatures (*e*.*g*., 37°C)^55^. Although at 37°C, particles have a high amount of thermal energy, causing rapid vibrations typically collapsing the wave function, coherence can persist if the quantum system is isolated from its environment such as trapped in a highly rigid molecular structure such as myelin wraps. The environment dynamically re-coheres the system faster than it decays if the observer’s measurement timeframes are extremely short (*e*.*g*., femtoseconds or nanoseconds)^55^.
- *No phonons*, there is no crystalline solid in proxy in which this quasi particle may vibrate.
- *No excitations in the surrounding material, no spontaneous electromagnetic emission, no fluctuating electromagnetic fields, no external environmental coupling*. the near-ideal topological quantum insulation of the intraperiod-myelin *V*_*IMI*_. system wrap, Fig. 19 a & b, up to 300 myelin wrap around one single axon and hundreds of identical myelin-wrapped axons in one nerve bundles provides further near-ideal topological insulation. It also supercharges the signal speed.
- *No electrode/insulator degrees of freedom, no degrees of freedom associated with the mechanism driving the ions, no charge-position fluctuations*. Ion channels *K*^*+*^/*Na*^*+*^ are pumping in their respective opposite direction, but in consecutive sequences^6,56.^
- The two ensembles of S_1_ and S_2_ of *Na*^*+*^, 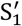 and 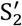 of *K*^*+*^, at each extremity of the cylinder, distance L between them, are separated at all times.

**Fig 18.**
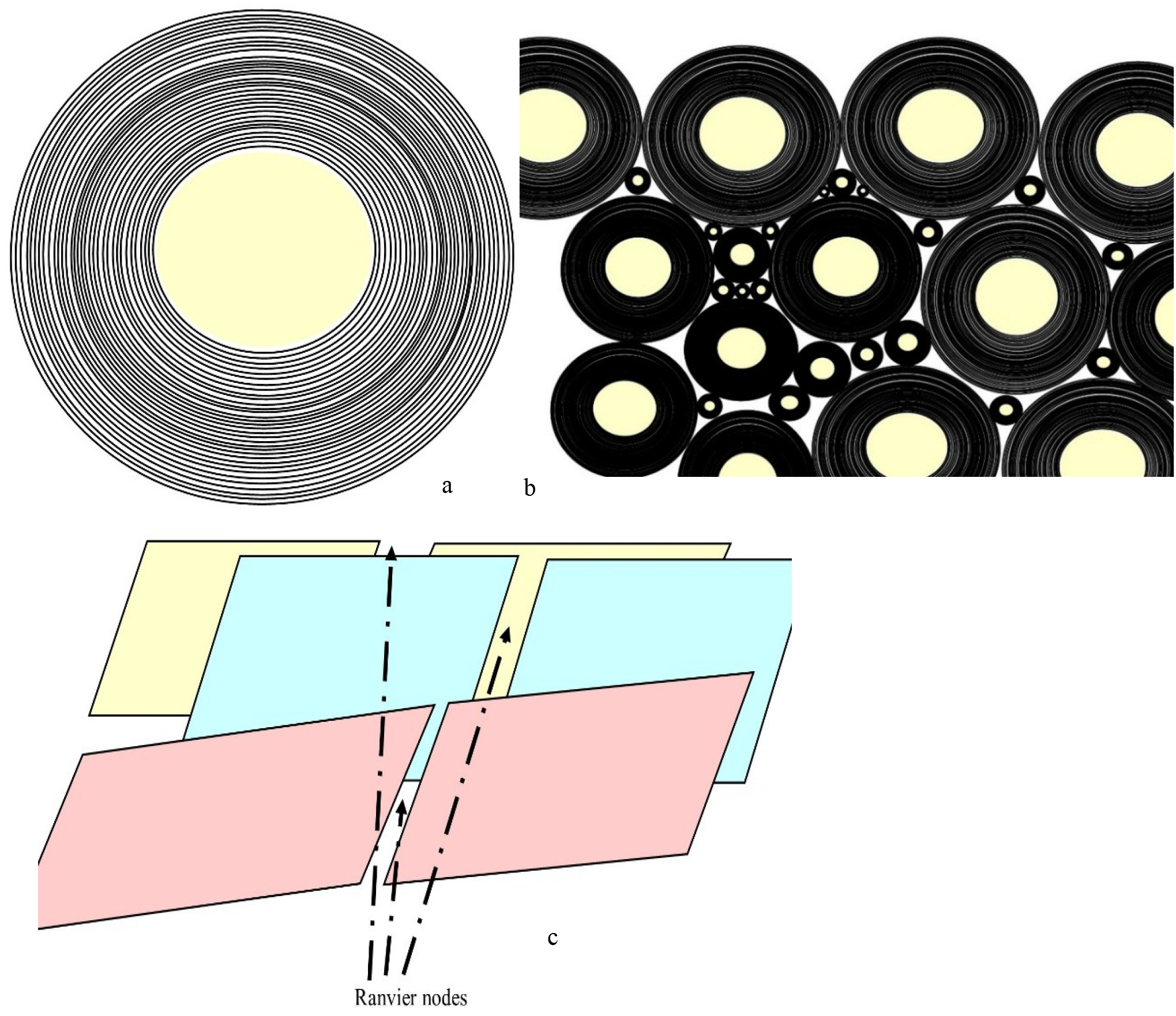
Myelin sheath. a). Transversal slice of the axon and its myelin insulation. Superposition of up to concentric 300 layers wrapping a single axon. Each one could host hundreds to thousands of niches, “memory cells”. b). Transversal slice of a long-distance white matter tract, the bundle may contain up to 200-200 million compressed myelinated axons. c). *2-D* plane of unfolded myelin layers exposing the random disjointed placement of Ranvier nodes exponentially increasing the hosting of charges ions in their memory cells. Axons within a single long white matter bundle fire asynchronously at different times.

**Fig 19.**
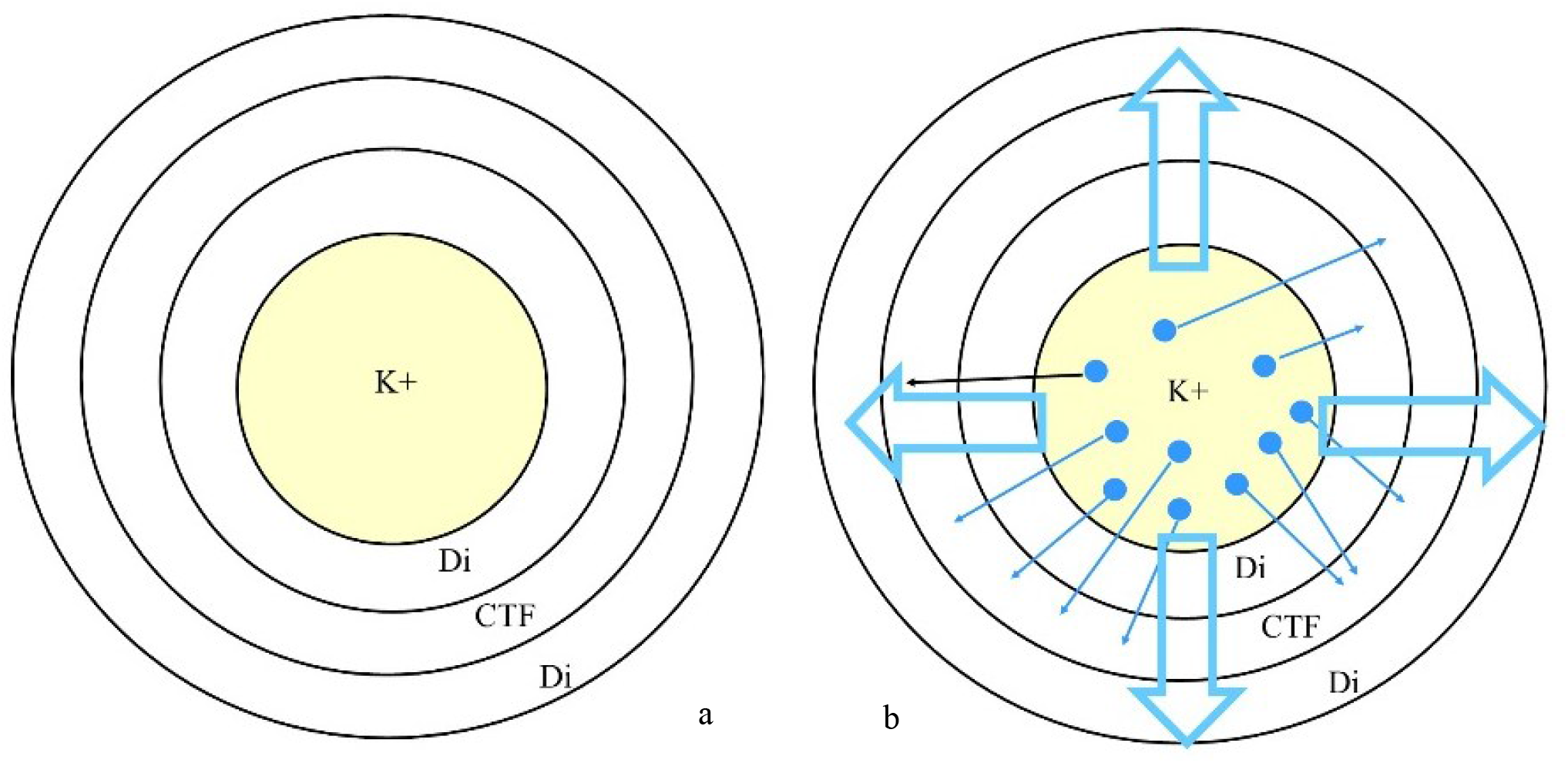
a). The axon and its concentric myelin multilayers restricted to two layers and a virtual space, “CTF”, charge trap flash, fused cell oligodendrocyte membranes. Di, dielectric, is a single myelin wrap. b). the action potential creates an electromagnetic field, then ion *K+* migrates in the extracellular milieu, and across the Ranvier nodes further into the virtual space between two myelin-dielectric layers. Some *K+* remaining trapped, and non-exposed, are storing nonvolatile memory in a CTF-like. The occurrence of a minimal action potential will lower the threshold and liberate the *K+* ions (as memory recall or stimulation). The data may be stored in 0 or 1. See text for details. Large transparent arrows with blue outline are the electromagnetic potential generated by AP. The trapping may occur in a myelin layer of the axon within or rather in adjacent myelin layers compressed in a bundle. See transversal section on Fig. 8 for details.

**Figure 20.**
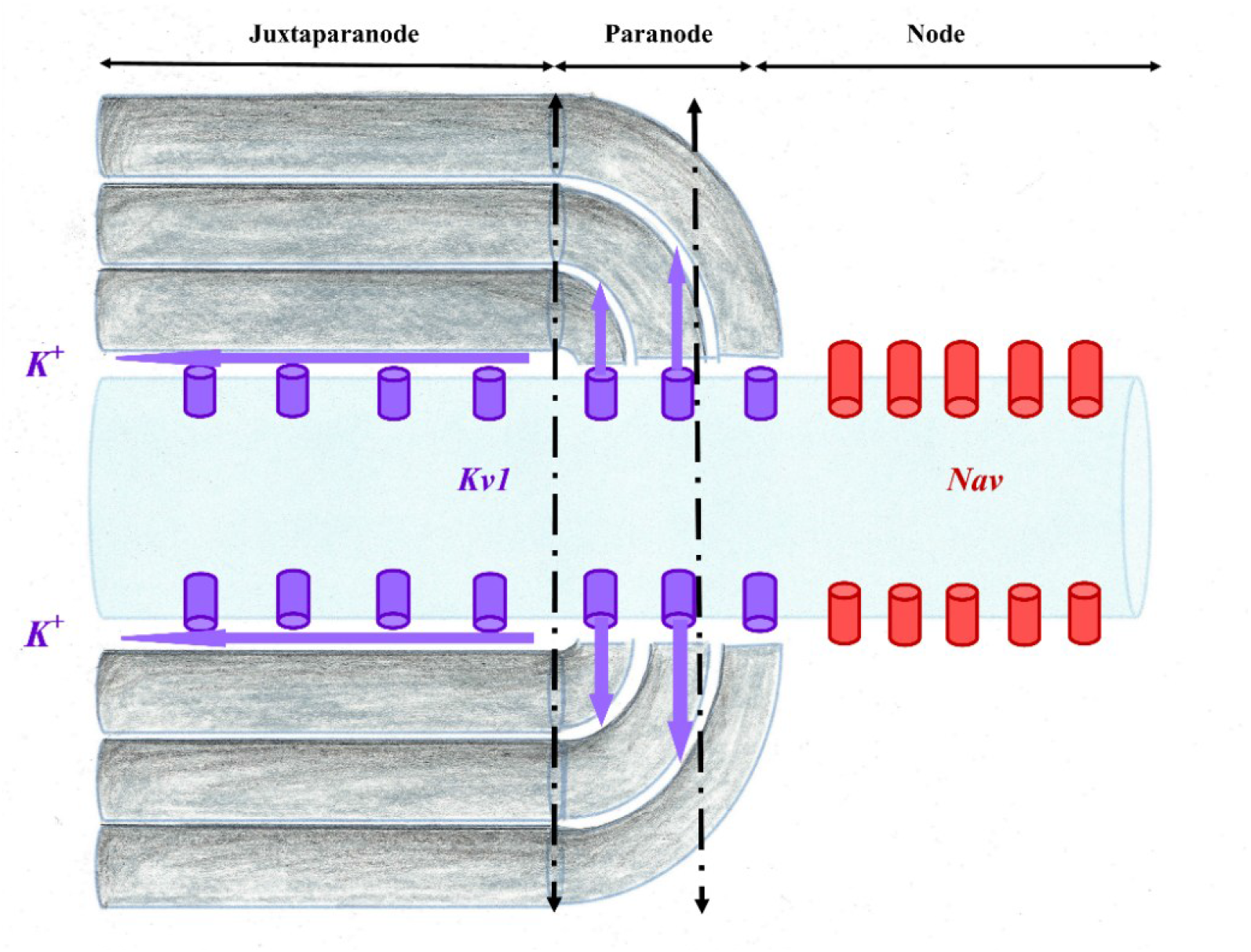
Node, paranode and juxtaparanode. *Nav* ion channels in red. in purple. Ions *K*^*+*^ may circulate within the axooligodendrial space (purple arrow) on the external surface of the myelin wrap. Storage of ion *K*^*+*^ occur externally and with two myelin wraps. Within a bundle, compressed myelin wraps are extremely hydrophobic, repelling water molecules and perfect insulators. From the outside in, the layers of myelin membrane compact, are bringing the inner and outer leaflets of consecutive lipid bilayers in extremely close apposition and extruding cytoplasm and extracellular fluid. In quantum mechanics, the space in-between can potentially be considered as within a topological insulator. This unique setting is likely preserving quantum coherence of the system. Once quantum tunneling occurred, ions *K*^*+*^ might be stored in the space between two myelin wraps. The overall system becoming potential CFT memory cell. Dotted lines: transversal sections are also expanded on a *2-D* plane in Fig. 7.

Influential physicists, including Penrose^57-59^, Hawking^57^, Wigner and others^60-63^ extensively devised on consciousness, brain, and quantum mechanics. Consciousness was not the focus of the current work, although considerations of brain function and quantum coherence were crucial.

#### i. Specific anatomy-histology preserving quantum coherence

The node of Ranvier is populated by voltage-gated *Na*^*+*^ channels, *Nav*, adjacent lies the paranode where the myelin is attached to the axon, next to the juxtaparanode where most voltage-gated *K*^*+*^ channels are located^64^. Voltage-gated *Na*^*+*^, *Nav*, channels, concentrated at the nodes, are separated from *K*^*+*^, *Kv1*, channels, clustered at the juxtaparanodal region by a specialized axooligodendroglial adhesion-contact that is formed between the axon and the myelinating cell at the paranode^65,66.^ Note that such *Nav* vs. *Kv1* separative property avoids potential decoherence by charged ions collisions. The paranode acts as a diffusion barrier which separates the *Nav* channels in the node of Ranvier from the *Kv1* channels in the juxtaparanodes (Fig. 8). The paranode “glues” the myelin to the axon, ensuring a tight connection between the myelin and the axolemma, and reducing current flow under the myelin sheath^64^. The strong adhesion between myelin and axolemma is achieved by cell adhesion molecules (see below)^64^. The juxtaparanode, adjacent to the paranode, where the axon is directly under compact myelin wraps, but does not extend for the entire length of the internode. Juxtaparanodes are clustered by Shaker-type *Kv1* channels.

The high density of *Nav* channels generates the maximum transmembrane voltage across the nodal membrane^67^. Any *Nav* channel misplaced to the internodal region wrapped with multiple myelin layers of myelin wraps, would become dysfunctional and generate a smaller voltage change. Most voltage-gated *K*^*+*^ channels, *Kv1*-type, are inserted in the juxtaparanodal region^64^. *K*^*+*^ channels, *Kv1* do not normally participate in generating AP. Blocking or knocking out *Kv1* channels did not affect propagation of PA^64^. Potential current flow is reduced by adhesive junctions between the ensheathing glial cells and axons^64^. These septate-type tight junctions leave a virtual space to the spiral pathway between the extracellular space at the node and the extracellular space at the juxtaparanode, through which some difference of potential (*K*^*+*^) may expand allowing partial activation of juxtaparanodal *K*^*+*^ channels. The exact level of AP activation depends on the current flow occurring under the myelin at the juxtaparanodal region, to maintain the extra-axonal voltage close to 0 mV. Structures associated with *Kv1* channels are found in the internodal and paranodal regions and are normally concealed under the myelin sheath^68^. The acceleration of myelin swelling after repetitive propagation of APs may be due to the opening-loosening of paranodal axooligodendroglial junctions^68^. From the outermost to the innermost layers, the myelin membrane undergoes progressive compaction, resulting in the close apposition of the inner tongue and outer leaflets of adjacent lipid bilayers. This process is extruding both cytoplasmic and extracellular fluid, thereby generating the highly compact structure characteristic of mature myelin^69^.

Nodal length and diameter are strictly regulated. The speed of nodal *Nav* channels activation depends on the capacitance of the node, and hence by its membrane area^64^. Pathologies increasing the area are slowing AP, unless extra *Nav* channels are inserted into the membrane to oppose this effect. Furthermore, an increase in node length, as also occurs in pathology, will increase the axial resistance for the current flow from the node to the internode that is needed to depolarize the next node, again slowing the action potential, or even causing it to fail^64^. Pathological disruption and retraction of the paranodal structure may loosen this junction^70^, ensuing preventing AP propagation, especially during repeated firing. Loss of internodal and paranodal adhesion results in overgrowth of nodes of Ranvier^71^. In multiple sclerosis, cerebral hypoperfusion and energy deprivation lead to elongation of the node of Ranvier, ensuing *Nav* channels covered with myelin wrap and disruption of paranodal junction, by loss of cell adhesion molecules (neurofascin 155, NF155), and paranodal contactin-associated protein (Caspr)^64^. This increases the virtual axooligodendroglial space between axolemma and myelin wrap, and *Nav* channels overlap with the paranodal protein Caspr^64^.

In addition, potassium channels *Kv1* within the juxtaparanodal axonal membrane may serve as depressing re-entrant excitation^72,73.^The astrocytic processes contain the inwardly rectifying *K+* channels *IRK1* and *IRK3*, which may allow *K*^*+*^ to accumulate during axonal activity^72,73.^

#### ii. Axooligodendroglial space and quantum coherence

The extracellular perinodal matrix is not a fluid or a liquid. Oligodendrocytes act as the primary producers of this specialized ECM medium. It takes the form of a dense, localized, gel-like macromolecular network that is structurally unique from the rest of the brain’s general interstitial matrix^74^. The molecular composition of the perinodal medium, rather than being fluid, liquid or generic connective tissue, this local medium is densely rich in specific structural and signaling macromolecules such as chondroitin sulfate proteoglycans, CSPG, highly negatively charged, hydration-retaining physical backbone of the node; brain link protein 1 anchors the proteoglycans into stable, higher-order aggregates; glycoproteins interacts directly with the axonal surface; hyaluronan the universal structural scaffold binding the proteoglycans^65,67.^The role of the perinodal matrix is ion buffering via the intense negative charge of the CSPGs creates a localized cation reservoir. This helps manage the rapid influx and efflux of *Na*^*+*^ and *K*^*+*^ ions required during saltatory electrical conduction^67,75.^It is also to molecular anchoring, It physically stabilizes and secures the clusters of voltage-gated sodium channels Nav on the exposed axon, keeping them from drifting into the myelinated segments^75^.

Axial conduction through the axooligodendroglial space is determining the saltatory AP propagation of action^69,76.^ Modulation of the axooligodendroglial space provides degrees of structural plasticity to the system^76^. Spatial learning alter the size of the axooligodendroglial space^76^. Modulating neuronal activity induces adaptive changes in myelin ultrastructure altering length of the node of Ranvier and width of the axooligodendroglial space (normal ≈ 12 nm), two of the axoglial parameters strongly determining AP conduction velocity, *V*. Furthermore, the inclusion or omission of axooligodendroglial space width could result in new internodes increasing or decreasing *V*, respectively^76^. Electrical conductance within the axooligodendroglial space facilitates the saltatory propagation of AP and changing the volume of this space theoretically alter *V*^76^. This property has implications for quantum tunneling of brain memory cells. Simulated *V* up to 3.5 times slower when the axooligodendroglial space width is set to 20 nm rather than 0 nm^76^.

#### iii. The unique system myelin intraperiod compartment isolating from microscopic environmental disturbances

Working memory training stimulates the production of myelinating oligodendrocytes from their precursor cells in the medial prefrontal cortex, anterior corpus callosum, dorsal thalamus and hippocampal formation^77^. Myelin sheath formation is also stimulated in the genu^77^. Myelin (white matter) is a highly hydrophobic structure composed of approximately 80% lipids, including galactocerebrosides and long-chain sphingomyelins, whose densely packed hydrocarbon chains confer strong water-repellent and electrical insulating properties. The remaining ∼20% consists primarily of structural proteins, proteolipid protein (PLP), among the most hydrophobic proteins known, whose transmembrane helices are deeply embedded within the lipid-rich membrane core. In the brain, myelin is proportionally thicker in thinner axons, measured as a *g*-ratio by electron microscopy^69^. The memory cells embedded within concentric 300 myelin layers wraps can be considered as dry solid-state relative vacuum (Fig. 8). Those virtual anatomical spaces of relative “negative or lower pressure points” are important for trapping in isolated niches of dry environment. Most biological mediums are made of fluids; however, specific anatomical 3D-organization of myelin wraps introduces localized conditions that are providing a dry, solid-state or vacuum-like conditions (relatively low-pressure environment) mitigating decoherence effects. This anatomical brain specificity makes it dry-hydrophobic with low-pressured air vacuum. The anatomical particularity of myelin wraps is that the *2-D* plane of unfolded myelin layers exposing the random disjointed placement of Ranvier nodes exponentially increasing the hosting of charges ions in their memory cells. This is also protecting the system myelin wrap-ion channel *K*^*+*^/*Na*^*+*^ from the fluidic environment and from firing, AP, because there are ∼300 myelin wraps around the axon in a bundle that may contain up to 200-200 million compress myelinated axons going from *2-D* to *3-D* or to a Hilbert Space. All of which, disjointing and isolating Ranvier nodes, only allowing very limited localization of transfers and AP propagation; this also warrants the precision of the signal in a bundle. Axons within a single long white matter bundle fire asynchronously at different times. Figure 6 illustrates the translation *3-D* to *2-D* allowing this asynchronous property and near-perfect isolation of the myelin wrap-ion channel *K*^*+*^/*Na*^*+*^ system allowing quantum coherence. It is randomly unlikely that two Ranvier nodes from two axons are facing each other. Hereby, given the myelin hydrophobic isolating the firing, rapid decoherence timescales 10^−13^ - 10^−20^ s within fluids will not affect firing timescales 10^−3^ - 10^−1^ s. Given the highly hydrophobic myelin wraps, collisions with water molecules is unlikely in the isolated myelin wrap-ion channel *K*^*+*^/*Na*^*+*^. In a bundle, myelin wraps are compressed and the oligodendrocyte intracellular space within the myelin wrap is likely virtual and squeezed enabling perfect insulation. Collisions with other ions is possible although unlikely since the general ion motion is mainly one-direction and because the cloud motion *K*^*+*^/*Na*^*+*^ in opposite direction occur at subsequent timeframes and in different locations *K*^*+*^ in paranode and juxtaparanode, *Na*^*+*^ in the node.

#### iv. Unique physical properties conferred by the node, paranode, and juxtaparanode

Ion channels *K*^*+*^/*Na*^*+*^ pumping (ATP-energy-system) driving ions against their electrochemical gradients, ***moving ionic charges, Na***^***+***^, ***K***^***+***^, in nodes, paranodes and juxtaparanodes, are creating an ***electromagnetic, EM, field***. It expresses a voltage, nodal intra- and extra-neuronal potential difference, which can be measured^27,56^,78, although action potential along the axon is ***not*** a conventional electric current relying on physical flowing of electrons through a conductor. The presence of the myelin sheath insulator, extracellular matrix and cells create a unique environment for propagating this EM field. Each ion channel *K*^*+*^/*Na*^*+*^ creates self-propagating electromagnetic waves jumping to the next ion channel, triggering the next wave, from next channel to next. This unique setting of myelin wrap-ion channel *K*^*+*^/*Na*^*+*^ systems, energy-dependent at each ion-channel, is an overall total energy-saving. Electromagnetic waves travel at the speed of light (299.10^6^ m/s) in perfect vacuum which is not the case across the extracellular matrix, or cytoplasm dropping the velocity down to 194.10^6^ to 214.10^6^ m/s. This EM wave velocity might be dropping through insulators like myelin sheath down to 124.10^6^-150.10^6^ m/s. This is approximately the propagation velocity of EM from node to node and start triggering the activation process of ion channels *K*^*+*^/*Na*^*+*^ pumping at the next node. Indeed, this is not the propagation velocity of AP, *V* ≈ 48 – 150 m/s. This drop of *V*, AP velocity, is time for ATP energy recruitment and activate the ion channels *K*^*+*^/*Na*^*+*^ pumping. The myelin sheath serves as an insulator for charged ions, from one node to next and as a dielectric in a node, paranode and juxtaparanode. Myelin might also behave as topological insulator for the EM waves, electrical insulator inside, but conduct electricity perfectly along the myelin-wrap surface, across extracellular matrix, or cytoplasm (Fig. 8). An EM wave triggers a new phase in distant charged ions primarily through mechanisms of EM-driven phase transitions^79,80^ such as for instance, by resonant energy transfer^81,82^ with potential phonon-phonon coupling^83,84^assuming a myelin-wrap may behave as topological insulator (*i*.*e*. “quantum material”), or by field-induced polarization^81,85.^When the oscillating electric field of an EM wave matches the natural frequency of a targeted ionic lattice or plasma, it drives collective structural transitions even across significant distances. Low-energy cyclotron emission is an established physical phenomenon. It occurs whenever any charged particle (like an electron) spirals in a magnetic field at low or non-relativistic energies^86^. The particle emits electromagnetic radiation at the cyclotron frequency, which is directly proportional to the magnetic field strength and inversely proportional to the particle’s mass.

Direct visualization of fast APs propagating along internodes remains infeasible due to EM propagation over a short internode length^4^. The dynamic phase-event takes several femtoseconds to propagate through a micrometer-scale internode; hence, it is too fast to be captured using existing phase imaging techniques relying primarily on binning temporal frames to improve sensitivity^4^. EM can propagate near the speed of light depending on the dielectric media^4^, such as myelin (124.10^6^-150.10^6^ m/s). To noninvasively, flexibly image the propagation of the fast transient excitation frame rates of up to 200 billion frames/s, a sequence depth of 350 frames, and the high phase sensitivity, differentially enhanced compressed ultrafast photography, Diff-CUP achieves noninvasive, direct spatiotemporal observation of passive internodal flows of electrical current, which plays a central role in AP propagation, and transmission of any other forms of electrical signaling across internodes where ion channel activities are absent. With a frame rate of up to 200 billion fps, a sequence depth of up to 350 frames, and a phase sensitivity of 20 μrad, Diff-CUP has enabled direct imaging of internodal current flows propagating in myelinated axons with an average conduction speed of 100m/s as well as the direct visualization of an EMP propagating in an LN crystal with a speed of 5 × 10^7^m/s. The reproduced internodal current flow speed and LN relative permittivity are consistent with the simulated and reported values, demonstrating the translation of our method into the biological and physical sciences. This team has reported compressed ultrafast photography (CUP), the world’s fastest camera that achieves both an ultrafast imaging of up to 70 trillion frames per second (fps) and a large sequence depth of up to 1000 frames^4^. However, without sufficient phase sensitivity (*e*.*g*., 0.9 mrad required to measure Aps) in spiking their previous CUP systems could not image either propagating APs or weak EM Pulses^4^. It is likely that an accurate assessment of the quantum physics underlying AP in the human brain will soon be made.

### D-III. The special case of Alzheimer’s disease provides further evidence for the proposed mechanism of memory storage

Oligodendrocyte and myelin disruptions are paramount factors in Alzheimer’s disease, AD, although the mechanisms remain unknown. Characteristic amyloid deposits, including spiral fibers around axons and dense helical coils at paranodal-juxtaparanodal loops, indicate amyloid infiltration within the axooligodendroglial space at paranodes, juxtaparanodes and internodes^87^. Additionally, we found aberrant myelination and disrupted paranodes on amyloid-plaque-associated dystrophic axons, a hallmark of AD known to impair action potential conduction and disrupt neuronal networks^87^. Despite no significant difference in the overall myelin coverage, there was greater localised paranode loss in AD brains compared with age-matched controls, indicating dysregulation of myelin-related signaling rather than a net loss of myelin. These results highlight significant paranodal protein alterations in AD, underscoring molecular disruptions at the axooligodendroglial space that may contribute to this pathogenesis^87^. Paranodal and periaxonal myelin regions are sites of amyloid-β accumulation^87^. This adds thickness and insulation within the axooligodendroglial space at paranodes, and juxtaparanodes, further blocking and preventing ions *K*^*+*^ to cross the axooligodendroglial space at paranodes, and juxtaparanodes. This would suggest that such thicker myelin wrap cannot serve as thin dielectric to achieve quantum tunneling and *K*^*+*^ cross out to memory cells within myelin wraps. Amyloid-β processing is dysregulated in the axooligodendroglial interface proteome and dense amyloid coils originated at paranodes and extended into juxtaparanodes^87^. The dense helical patterns likely represent amyloid-β accumulation within paranodal channels, while the less dense spirals may be extensions into periaxonal spaces at internodes. Amyloid spiral and coil around an axon within the axooligodendroglial space are localized at paranodes and juxtaparanodes^87^. This invasive axooligodendroglial pathological process may prevent normal quantum tunneling into thicker dielectric myelin layer. Bulbous axonal enlargements were sometimes observed at these helical amyloid-β aggregation sites. All of which occur without overt myelin loss^87^, rendering difficult the imaging diagnosis in AD patients.

## E. Conclusion

The myelin layers-intraperiod *V*_*IMI*_ compartment system, constitutes a unique biological near-ideal topological insulation in the human brain, and in the gnathosthomata (including familiar classes such as mammals) that possess both a centralized brain and true myelin sheaths. We tested stringent biological and physical boundaries which must be exact because a single missing/inexact element defining the anatomy, physiology, biochemistry, cytology, histology, and nano, leads to inaccurate quantum equations or interpretations of the myelin system. The equations of quantum mechanics are exact, and are exceptionally accurate predictors that are well-known to pass any further single experimental test. A rigorous inventory of the insulation of the intraperiod *V*_*IMI*_ void from external factors and the potential channels of decoherence led us to identify a threshold of quantum coherence to decoherence. Therefore, propagating internodal electromagnetic interaction through the *V*_*IMI*_ void can be represented via Hamiltonian. Furthermore, under higher restricted parameter threshold, entanglement was achievable.

Our next logical step was to investigate three potential mechanisms for ion *K*^*+*^ data storage. Preserving a near-topological quantum insulation limits the possibilities, and reduced the options to three hypotheses 1. Quantum tunneling through a dielectric, *i*.*e*. single myelin layer. 2. Quantum tunneling through squeezing void, *V*_*IMI*_. The action potential is accompanied by mechanical axonal waves and myelin layers compression. Compression reduces the WKB action, and exponentially increase in transmission-tunneling probability. Resonance tunneling may add orders-of-magnitude boosts. 3. Classical squeezing passage.

Quantum tunneling (Model 1) across a single myelin layer under highly restricted parameters. We could only assess that option is a possibility, although an exact evaluation of how many ions *K*^*+*^ may be trapped in the charge-trap, intraperiod *V*_*IMI*_, was not possible. Some parameters at the quantum scale (nm, Å) were not yet available. The size and weight of the ion *K+* was also a limiting factor compared to the electron.

The second hypothesis (Model 2) through the intraperiod *V*_*IMI*_ virtual void into the charge-trap, intraperiod *V*_*IMI*_. The charge-trap, intraperiod *V*_*IMI*_, has two purposes, a). Dielectric opening to the ion *K+*; and b). Charge-trap, capturing and storing the ion *K+*. This option seemed the most easily attainable, and plausible, under less restricted parameters. Resonant tunneling provided several orders-of-magnitude boosts. The synergy of resonant tunneling, compression, final gap, barrier height, electric field, was boosting the likelihood of quantum tunneling to a probability of *P* ≈ 0.4 − 1.

Unlike quantum tunneling, classical physic passage does not allow fine voltage-controlled metering, sorting, organizing the data storage, later rendering later retrieving impossible. In addition, we were unable to assess what local force would become sufficient to open the intraperiod *V*_*IMI*_ virtual void or cross the single myelin layer. Therefore, we were unable to test this option.

Hypotheses 1 and 2 are totally in line with earlier findings about memory storage, LTP, Hebbian theory, and synapse strengthening^1,27-30^,49,88. This is also in agreement with past studies on Alzheimer’s disease and other neurodegenerative studies, further shedding light on their pathological mechanisms.

## Acknowledgements

Patents pending (**63/984**,**445 & 64/175**,**133**), Dr. Philip P Foster, filed in 2026 with *USPTO/EPO*. This work was supported partly by a *National Science Foundation* Grant # *CBET* **2224942** to Dr. Aladin M Boriek. We are thankful to Dr. Roland Glowinski, Department of Mathematics & Statistics, University of Houston & University Pierre & Marie Curie, Paris, France. We dedicated this work to Dr. Roland Glowinski.

